# Targeting mitochondria in cancer therapy could provide a basis for the selective anti-cancer activity

**DOI:** 10.1101/432252

**Authors:** Dmitri Rozanov, Anton Cheltsov, Aaron Nilsen, Christopher Boniface, James Korkola, Joe Gray, Jeffrey Tyner, Cristina E. Tognon, Gordon B. Mills, Paul Spellman

## Abstract

To determine the target of the recently identified lead compound NSC130362 that is responsible for its selective anti-cancer efficacy and safety in normal cells, structure-activity relationship (SAR) studies were conducted. First, NSC13062 was validated as a starting compound for the described SAR studies in a variety of cell-based viability assays. Then, a small library of 1,4-naphthoquinines (1,4-NQs) and quinoline-5,8-diones was tested in cell viability assays using pancreatic cancer MIA PaCa-2 cells and normal human hepatocytes. The obtained data allowed us to select a set of both non-toxic compounds that preferentially induced apoptosis in cancer cells and toxic compounds that induced apoptosis in both cancer and normal cells. Anti-cancer activity of the selected non-toxic compounds was confirmed in viability assays using breast cancer HCC1187 cells. Consequently, the two sets of compounds were tested in multiple cell-based and *in vitro* activity assays to identify key factors responsible for the observed activity. Inhibition of the mitochondrial electron transfer chain (ETC) is a key distinguishing activity between the non-toxic and toxic compounds. Finally, we developed a mathematical model that was able to distinguish these two sets of compounds. The development of this model supports our conclusion that appropriate quantitative SAR (QSAR) models have the potential to be employed to develop anticancer compounds with improved potency while maintaining non-toxicity to normal cells.

## Materials and Methods

### Reagents

All reagents were from Sigma, unless otherwise indicated. CellTiter-Glo reagent was from Promega. Glutathione reductase (GSR) activity kit was from Cayman. GSR producing plasmid was a kind gift of Dr. Becker (Justus-Liebig University Giessen). GSR was expressed in BL21(DE3) cells and purified by both metal chelating and affinity chromatography on 2’,5’-ADP-Sepharose as described [1].

### Cells

Human prostate carcinoma, breast, and pancreatic carcinoma cells were obtained from ATCC. Chemotherapy resistant prostate carcinoma cells were from Dr. Korkola. Human primary hepatocytes were obtained from Lonza. All cells were cultured according to the provider’s guidelines. Bone marrow aspirates or peripheral blood samples were collected from acute myeloid leukemia (AML) patients under an OHSU Institutional Review Board (IRB) approved research collection protocol which covers *in vitro* drug testing of leukemia cells and genetic studies. Patients signed an IRB-approved written consent form after verbal consent was obtained. Mononuclear cells were isolated from peripheral blood samples using a Ficoll gradient and viability of isolated mononuclear cells was assayed using Guava Viacount reagent. All patients were treated in accordance with the ethical guidelines laid out in the Declaration of Helsinki. AML cells were cultured in RPMI medium supplemented with 10% fetal bovine serum.

### Hydrogen peroxide cell-based assay kit

Cells were incubated for 2 h with either compounds (30 μM) or DMSO in a 96-well plate. After the treatment, 80 μl of cell culture medium was collected and the level of hydrogen peroxide was determined using a Hydrogen Peroxide Cell-Based Assay kit (Cayman).

### Glutathione reductase (GSR) activity assay

Recombinant GSR (0.5 u/well) in a 96-well plate was incubated in the GSR assay buffer (50 mM potassium phosphate buffer, pH 7.5 supplemented with 1 mM EDTA), 100 μl with and without compound (30 μM) for 2 h at 25°C. At the end of incubation, 100 μl of solution containing NADPH (0.087 mM), oxidized glutathione (GSSG) (0.95 mM) and 1 μl of the reagent that detects ratio of GSSG to reduced glutathione (GSH) (Abcam) were added and the mixture was incubated for an additional 1 h at 25°C. GSR activity was measured by using 490-nm excitation and monitoring fluorescence at 520 nm with a fluorescence plate reader. One unit of GSR activity was defined as the amount of enzyme required to produce 1 μM of GSH in a total reaction volume of 200 μl in 1 minute in the GSR assay buffer containing NADPH (0.087 mM), GSSG (0.95 mM) and 1 μl of GSH/GSSG ratio detection reagent (Abcam).

### NADH/NADPH dehydrogenase-mediated ROS generation assay

Recombinant NADH/NADPH dehydrogenase (0.5 units/well) in a 96-well plate was incubated in 100 μl of the assay buffer (50 mM potassium phosphate buffer, pH 7.5 supplemented with 1 mM EDTA) containing NADPH (3 μM) and superoxide dismutase (SOD) (100 units/well) with and without compound (30 μM) for 4 h at 37°C. At the end of incubation, the level of hydrogen peroxide was measured using a Hydrogen Peroxide Assay kit (Cayman). Control samples were without NADH/NADPH dehydrogenase. The rate of NADH/NADPH dehydrogenase- and compound-mediated ROS production was determined by subtracting values that were obtained in the absence of NADPH dehydrogenase. One unit of NADH/NADPH dehydrogenase was defined as the amount of enzyme required to reduce 1.0 μmole cytochrome C per min/mg in the presence of menadione substrate at 37°C. One unit of SOD was defined as the amount of enzyme required to inhibit reduction of cytochrome C by 50% in a coupled system with xanthine oxidase at pH 7.8 at 25°C in a 3.0 mL reaction volume. Xanthine oxidase concentration should produce an initial ΔA_550_ of 0.025 ± 0.005 per min.

### MitoCheck Complex I activity assay

MitoCheck Complex I activity was determined as described in the manual of the manufacturer’s manual (Cayman). Briefly, compounds (30 μM) or DMSO were mixed with the Complex I activity assay buffer that contained bovine heart mitochondria reagent, NADH, and ubiquinone in the presence of KCN to inhibit complex IV activity. The level of Complex I activity was monitored in a kinetic assay of NADH oxidation by measuring a decrease in absorbance at 340 nm. The residual Complex I activity was calculated by dividing the rate of compound wells by the rate of DMSO wells for the linear portion of the curve.

### Glyceraldehyde-3-phosphate dehydrogenase (GAPDH) activity assay

Cells were incubated for 2 h with either compounds (30 μM) or DMSO in a 6-well plate. Next, cells were collected in 200 μl of PBS and disrupted by sonication. Insoluble material was removed by centrifugation. The level of GAPDH activity was determined by GAPDH activity assay kit (BioVision).

### Small-molecule compounds

NSC130362 and its analogs were obtained from the NCI/DTP Open Chemical Repository (http://dtp.nci.nih.gov/repositories.html). Clinical drugs were purchased as dry powders from Cayman.

### Statistical analysis

The results of the different assays are expressed as mean values based on at least three replicates. The statistical significance of differences in the treatment outcomes was determined by one-way ANOVA or Mann-Whitney U tests.

### Introduction

Despite the advances achieved in the detection and treatment of early cancer that have contributed to declining cancer-specific mortality in the United States, metastatic cancer remains in most cases an incurable disease. In this context, identifying new drugs and designing more efficacious and safe cancer treatments to prevent relapse in patients and to treat metastatic disease are clearly needed to provide an impact on cancer mortality rates.

One promising strategy for successful cancer therapy is to induce oxidative stress and followed by apoptosis in cancer cells but not in normal cells. Elevated levels of reactive oxygen species (ROS) and subsequent oxidative stress are hallmarks of carcinogenesis and metastasis providing a potential therapeutic index [2–4]. Our data and recent studies by others demonstrated that elevated levels of ROS can be exploited *in vitro* and *in vivo* to preferentially target cancer cells while sparing normal cells [5–8]. The ROS-based approach to induce apoptosis in cancer cells is conceptionally different from conventional therapy targeting well known oncogenes and tumor suppressors - a therapy which is often ineffective due to multiple genetic and epigenetic alterations in cancer cells and the ability of cancer cells to upregulate compensatory mechanisms [9, 10]. The shortcomings of conventional targeted therapy approaches have prompted the development of alternative approaches. Instead of targeting specific oncogenes and tumor suppressors, exploiting common biochemical alterations in cancer cells, such as an increased ROS stress, could provide the basis for developing selective and potent therapeutic agents.

To cope with increased production of ROS, mammalian cells have developed two major electron donor systems, the thioredoxin (Trx) system and the glutathione (GSH) system [11, 12]. The Trx redox system is composed of thioredoxin reductase (TrxR), Trx, and NADPH while the GSH redox system is composed of GSR, GSH, and NADPH. The Trx and GSH system represent two complementary defense systems against oxidative stress. Other redox-sensitive enzymes that play a role in the oxidative stress response include Trx- and GSH-peroxidase, GSH- S-transferase (GST), and isocitrate dehydrogenase [13–15]. Thus, targeting any of these components can potentially induce oxidative stress which can result in cell death.

We recently reported the discovery of 1,4-naphthoquinine (1,4-NQ) derivative, NSC130362, which inhibits GSR and, as a consequence, induces oxidative stress and subsequent apoptosis in cancer cells but not in normal human primary hepatocytes. NSC130362 also showed anti-tumor activity *in vivo* [8]. In addition to inhibiting GSR, 1,4- NQs can be reduced by NADH/NADPH dehydrogenase followed by autoxidation, which results in the formation of ROS and potential oxidative stress. The extent of autoxidation is dependent on the type and position of substituents. 1,4-NQs can also reduce cell viability *via* arylation of cellular nucleophiles such as GSH, DNA, RNA and proteins and also by inhibition of DNA synthesis or mitochondrial function [16–18]. In the current work, we tested different activities of NSC130362 and its analogs with the aim of identifying the factors responsible for enabling NSC130362’s selective anti-tumor activity. Based on the obtained results, we were able to construct a mathematical model that could distinguish toxic NSC130362 analogs from analogs that were non-toxic to normal cells. The goal of our study was to demonstrate that it is possible to build a quantitative structure-activity relationship (QSAR) model that can guide optimization of potent and non-toxic compounds.

## Results

### Characterization of NSC130362 as a lead compound

The NIH has defined several requirements that should be met for a compound to be considered as a starting material for drug discovery programs [19]. Starting compounds should [1] elicit a reproducible response in at least two assay types and have a dose-response over a hundred-fold concentration range; [2] be analytically validated in terms of integrity and purity; [3] demonstrate adequate potency; [4] possess a tractable starting point of chemical optimization with no obvious major chemical liabilities. Most of these requirements were addressed in our previous publication [8]. Thus, we demonstrated that our lead compound, NSC130362 (Fig 1, A) selectively induces reproducible responses in either oxidative stress or caspase 3/7 activity assays in cancer cells. We also showed that its combination with different oxidative stress inducers, such as arsenic trioxide (ATO), myricetin (Myr), and buthionine sulfoximine (BSO), caused cell death in a variety of breast, pancreatic, prostate, and lung carcinoma cell lines as well as in human melanoma MDA-MB-435 and AML cells from patients. Finally, NSC130362 demonstrated anti-tumor activity *in vivo* [8]. To complement these results, we performed dose-response studies in pancreatic carcinoma MIA PaCa-2 cells (Fig 1, B). The obtained data clearly showed that NSC130362 induced a dose-response over a hundred-fold concentration range in the presence of ATO. In addition, the structure of NSC130362 was verified *via* proton NMR and compound purity was at least 95%. According to the structure, this compound can be easily modified to produce chemically related analogs [20].

**Fig 1.**
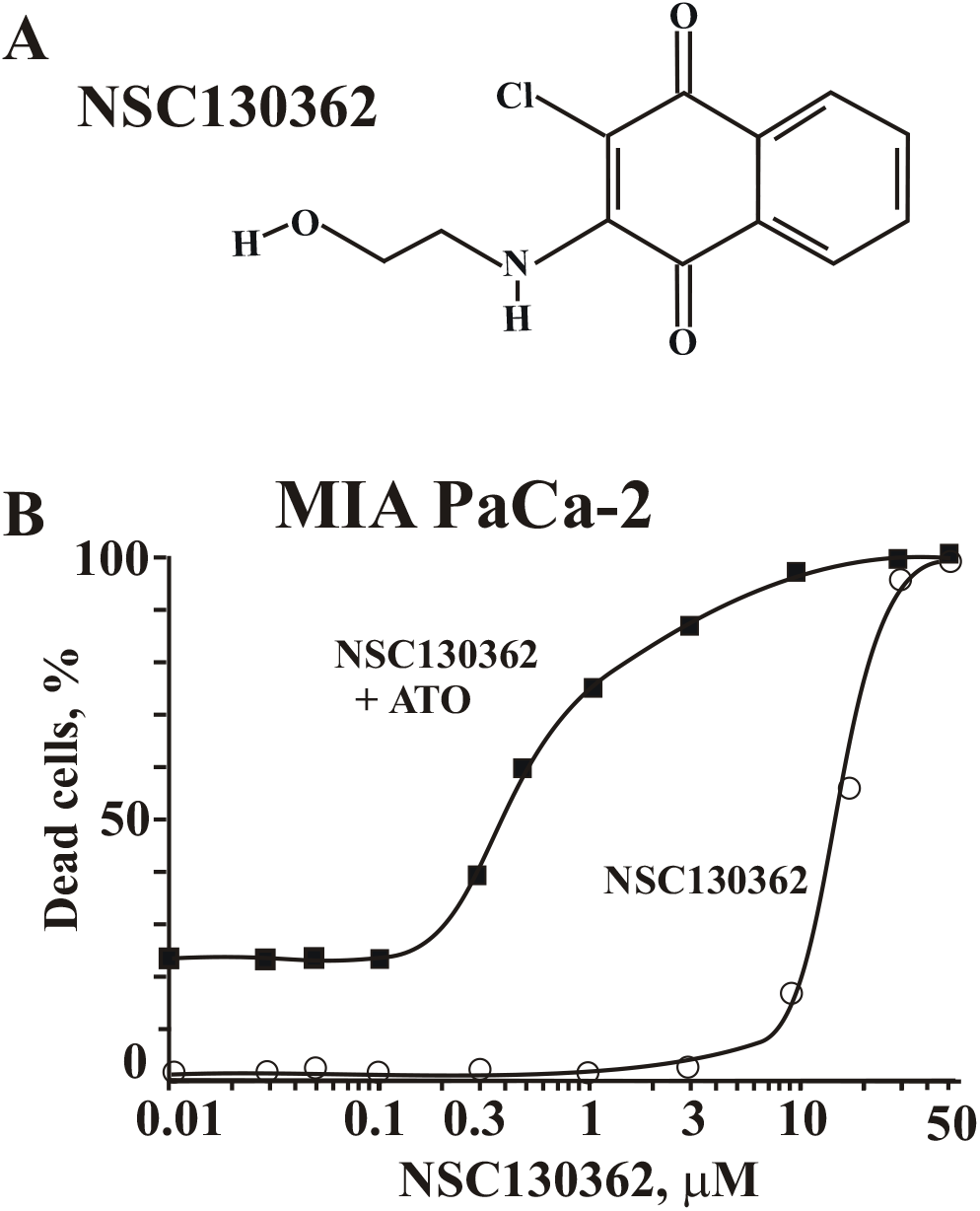
A, Structure of NSC130362. B, Dose response curve of NSC130362 in MIA PaCa-2 cells. Cells were preincubated with NSC130362 for 2 h followed by addition of ATO (5 μM) and incubation for an additional 24 h. At the end of the treatment, the ratio of dead cells was determined by a CellTiterGlo reagent. *P*<0.05. NSC130362 is a 1,4- NQ. Quinone moieties are present in many drugs such as anthracyclines, daunorubicin, doxorubicin, mitomycin, and saintopin, which are used in anti-cancer therapy [16]. The effects of quinones on cell viability are mainly due to generation of reactive oxygen species (ROS) or to covalent attachment to biomolecules *via* arylation reactions [16, 21]. In addition to their ability to induce oxidative stress, ROS-generating agents can also potentiate the ability of clinical drugs to induce apoptosis in cancer cells [22–24]. To test whether NSC130362 is able to promote the cytotoxic effect of anti-cancer drugs, we performed cell viability assays using chemotherapy resistant PANC1 and YAPC pancreatic carcinoma cells (Fig 2). We found that NSC130362 significantly increased the ability of all tested pancreatic cancer drugs [25] to induce apoptosis in PANC1 and YAPC cell lines. To further validate these findings, we analyzed a variety of chemotherapy resistant prostate carcinoma cells in cell viability assays using prostate cancer drugs docetaxel and enzalutamide (MDV3100) either alone or in combination with NSC130362 (Fig 3). For comparison purposes, we also included FDA-approved ATO in this analysis. Our selection of ATO was based on our previous findings showing that ATO and NSC130362 exhibited synergistic responses against multiple cancer cells [8]. As we expected, NSC130362 efficiently potentiated the effect on cell viability of either ATO or MDV3100. NSC130362 also potentiated docetaxel-induced cytotoxicity, but to a lesser extent most likely because docetaxel was more cytotoxic as monotherapy than MDV3100 and ATO. Finally, we performed screening of small-molecule kinase inhibitor libraries [26] with and without NSC130362 against freshly isolated primary acute myeloid leukemia (AML) cells obtained from patients. Encouragingly, NSC130362 decreased EC50 (cytotoxicity) of several kinase inhibitors by 1000-fold (Supporting Information Table S1). We also found that there were a number of shared kinase inhibitors across leukemia cells from three different patients whose activity was significantly induced (more than 10 times) by NSC130362. In summary, in these independent analyses, NSC130362 exhibited synergistic anti-tumor activity against different cancer cell types when combined with distinct FDA-approved drugs and kinase inhibitors. Based on these findings, we conclude that NSC139362 is a promising starting compound for further characterization.

**Fig 2.**
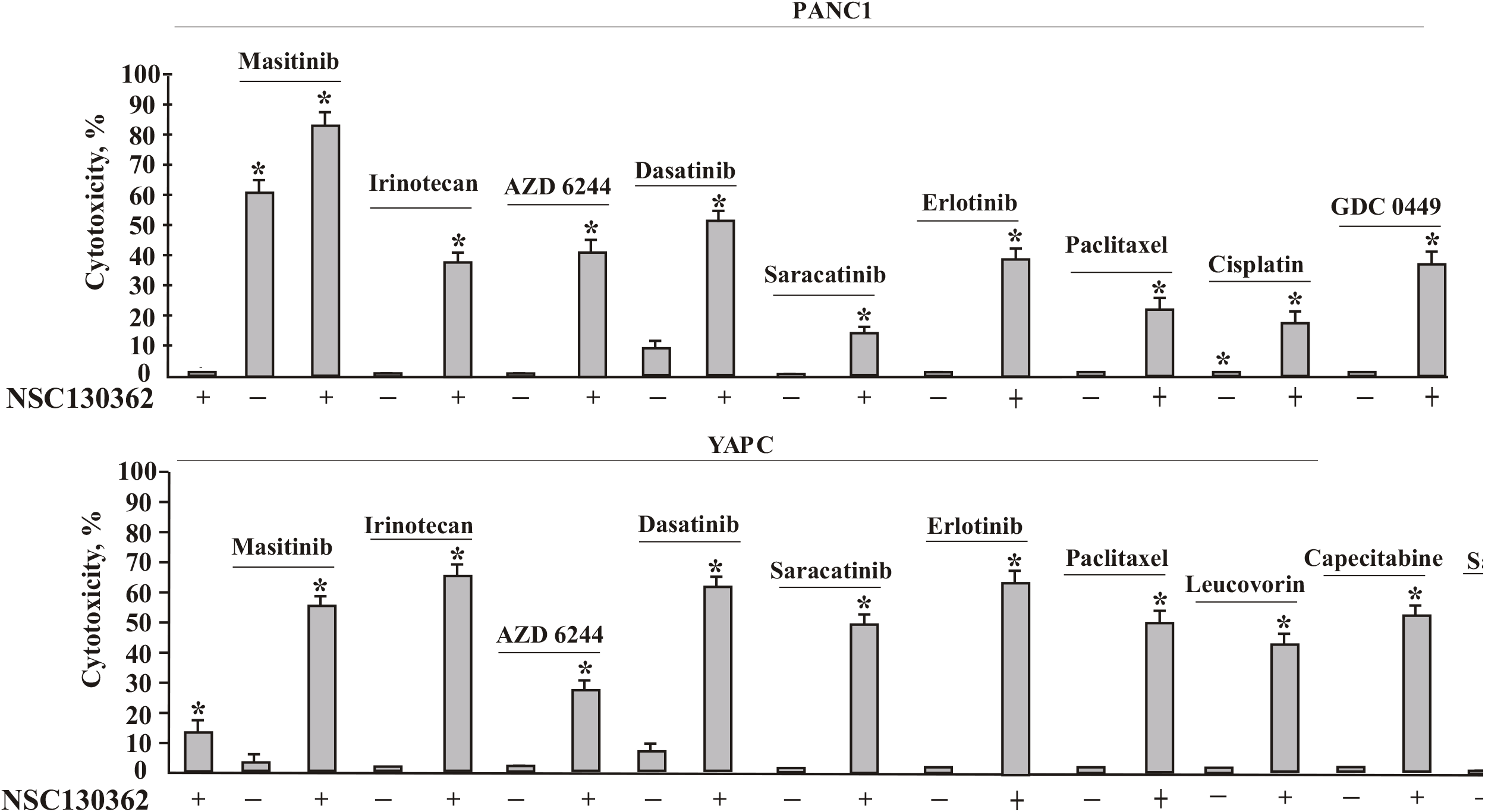
Combined treatment of NSC130362 with pancreatic cancer drugs against PANC1 and YAPC cells. Cells were pre-incubated with NSC130362 (10 μM) for 2 h followed by addition of drugs (10 μM) and incubation for an additional 24 h. At the end of the treatment, the ratio of dead cells was determined by a CellTiterGlo reagent. The first bar is NSC130362 alone. *, *P*<0.05.

**Fig 3.**
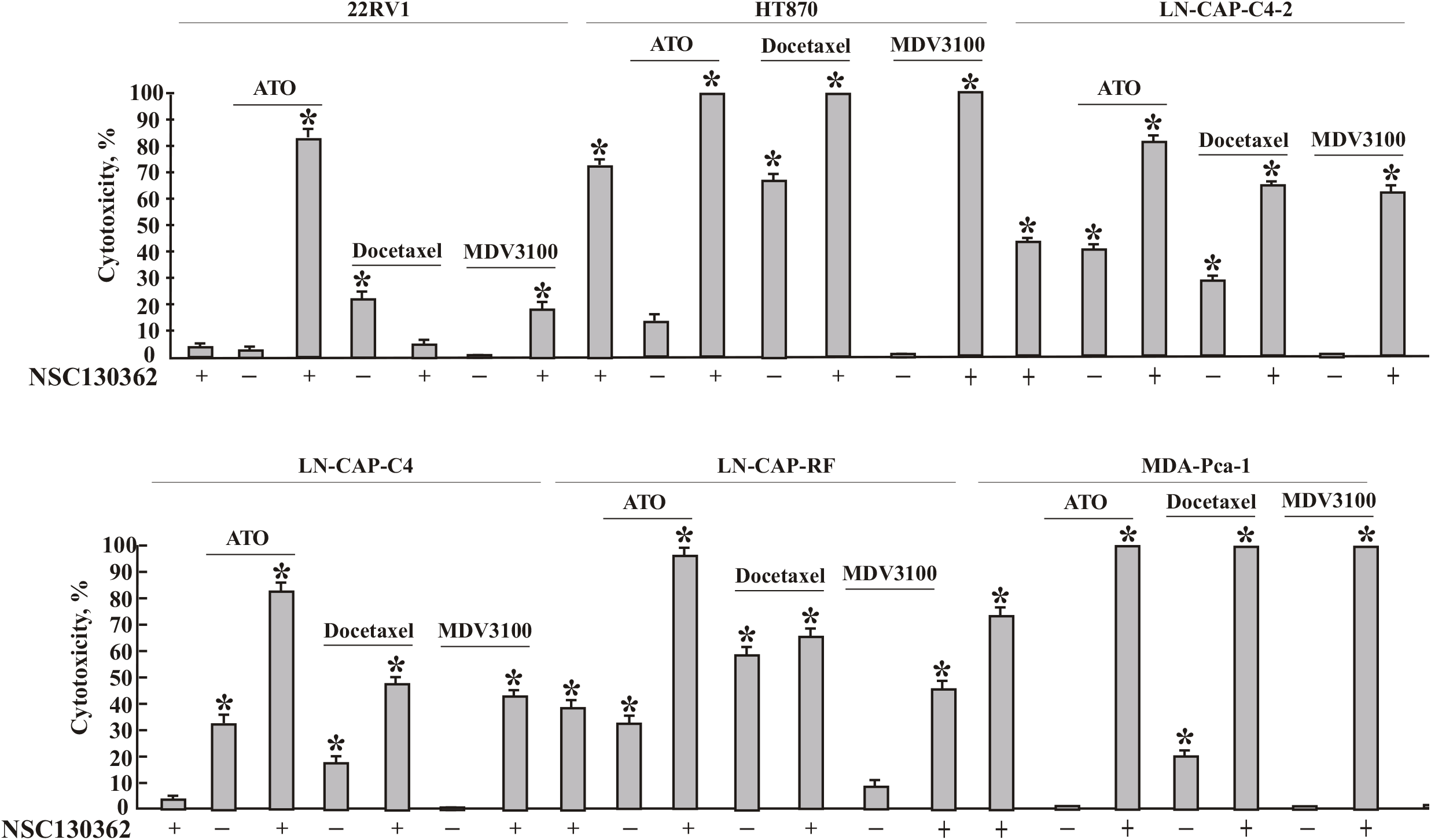
Combined treatment of NSC130362 with ATO, docetaxel, and MDV3100 against prostate cancer cells. Cells were pre-incubated with NSC130362 (10 μM) for 2 h followed by addition of drugs (10 μM) and incubation for an additional 24 h. At the end of the treatment, the ratio of dead cells was determined by a CellTiterGlo reagent. . The first bar in the treatment of each cell line is NSC130362 alone. *, *P*<0.05.

### Mechanisms underlying NSC130362-mediated effects in cancer cells

To elucidate the mechanism of synergy between NSC13062 and ATO, we analyzed the effect of a reducing environment on cytotoxic activity mediated by the NSC130362/ATO combination. The rationale for this assay was based on the knowledge that NSC130362, in addition to its ability to induce ROS *via* NADH/NADPH dehydrogenase-mediated reaction, targets GSR, an enzyme that maintains the reducing environment of the cell and is critical in resisting oxidative stress. In turn, ATO interferes with numerous intracellular signal transduction pathways that may result in the induction of apoptosis, the inhibition of tumor growth, and angiogenesis [27]. These effects, at least in part, are mediated *via* interaction of ATO with redox-sensitive proteins and enzymes. Almost all ATO-interacting proteins are protected against oxidation by the highly reduced intracellular environment that is controlled by GSR. Thus, GSR inhibition could facilitate ATO-mediated oxidation of intracellular proteins resulting in a synergistic induction of apoptosis in cancer cells. In agreement, our data showed that the effect of NSC130362/ATO combination is inhibited by dithiothreitol (DTT) (Fig 4). Thus, we conclude that the effect of NSC130362 on ATO-mediated cytotoxicity is mediated, at least in part, by its ability to induce ROS and/or inhibit GSR. We also tested the effect of hypoxic conditions on the activity of NSC130362/ATO combination. The effect of low level of oxygen on NSC130362/ATO-mediated cytotoxicity in MIA PaCa-2 cells was less pronounced than that of DTT (Fig 4).

**Fig 4.**
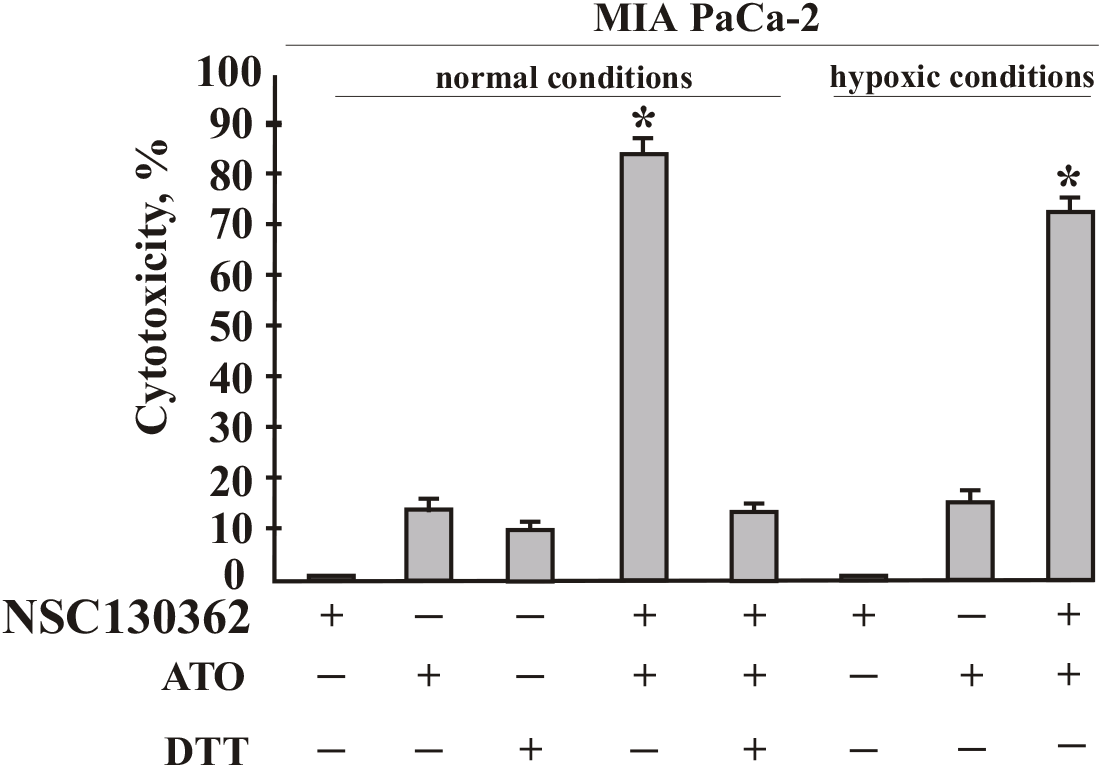
Effect of DTT and hypoxic conditions on the activity of NSC130362/ATO combination in MIA PaCa-2 cells. Cells were pre-incubated with NSC130362 (3 μM) in either normal (24% O_2_) or hypoxic (0.5% O_2_) conditions for 2 h in the presence or absence of DTT (3 mM) followed by addition of ATO (5 μM) and incubation for an additional 24 h. At the end of the treatment, the ratio of dead cells was determined by a CellTiterGlo reagent. *P*<0.05.

### Structure-activity relationship studies of NSC130362 analogs

To evaluate the effect of substituents on the cytotoxic activity of NSC130362 analogs toward cancer and normal cells, we analyzed cell viability after treatment with compounds in the absence or presence of ATO using pancreatic cancer MIA PaCa-2 cells and human primary hepatocytes. We selected MIA PaCa-2 cells because these cells are highly responsive to ROS [8]. These cells were also selected because of their robust engraftment in immunodeficient mice and sensitivity to ROS-inducing agents *in vivo.* Importantly, the results obtained in MIA PaCa-2 cells are similar to those obtained in breast and other cancer cells [8]. We selected hepatocytes as normal cells because hepatocytes are mediate toxicity to multiple drugs and are used as a standard to assess toxicity of drugs *in vitro* [28]. We then determined the safety index of each NSC130362 analog by dividing the EC50 (cytotoxicity) measured in hepatocytes by that obtained in cancer cells in the presence of ATO. Because ATO and NSC130362 showed clear synergy against MIA PACa-2 cells (Fig 1) [8], the safety index was related to the combined treatment, however, data for monotherapy were also provided. ATO at the concentration of 5 μM did not affect the viability of both MIA PACa-2 cells and human hepatocytes. For comparison with the standard of care, doxorubicin was also included in this analysis. In addition to its effect on DNA replication, doxorubicin is also known as an oxidative stress inducer [29].

We performed SAR studies to understand the chemical-biological interaction controlling cytotoxicity in MIA PaCa-2 cells without affecting the viability of human hepatocytes. To cover more chemical space, in addition to 1,4-NQs, quinoline-5,8-diones were also analyzed (Fig 5).

**Fig 5.**
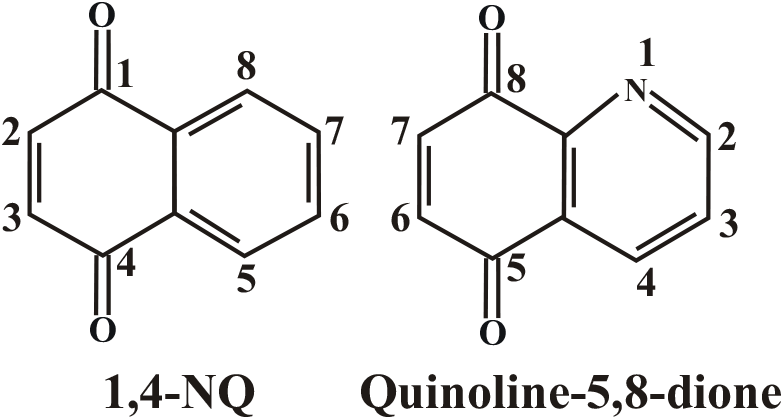
Structure of 1,4-NQ and quinoline-5,8-dione.

First, we analyzed 2-chloro-3-amino-1,4-NQ and 2-chloro-3-(alkylamino)-1,4-NQs (Table 1). SAR results showed that the cytotoxic activity of compounds **1–9** in cancer cells depends on the size of the 3-substituent. The anti-cancer potency and safety index decreased with increased length of the substituent. When compounds with branched 3-(alkylamino) substituent (compounds **10–13**) were analyzed, we also noticed a decrease in cytotoxic activity and safety index when the length of alkyl chain is changed from isopropyl (compounds **10**) to either sec-butyl or isobutyl (compounds **11** and **12**, respectively). However, when the alkyl group was tert-butyl (compounds **13**) there was a significant increase in the anti-cancer activity and safety index. We conclude that the shape of 4-carbon alkyl group plays an important role in the biological activity of the analyzed compounds. The importance of the shape of the alkyl substituent in 2-chloro-3-(alkylamino)-1,4-NQs can be drawn from study of compounds **14–16**. In this case, when two hydrogens of the amino group are substituted by either two methyl groups (compound **14**) or aziridine groups (compound **15**), the resulting compounds have similar cytotoxic activity in MIA PaCa-2 cells to a compound when one hydrogen of the amino group is substituted by the methyl group (compound **2**). However, the effect of these substitutions on toxicity to hepatocytes was more pronounced. Compounds **14** and **15** were approximately 2 and 6.5 times more toxic toward hepatocytes than compound **2**, respectively. Strikingly, when two methyl groups were attached to the aziridine group (compound **16**), the compound became completely non-toxic against normal cells, while maintaining almost identical levels of anticancer activity with that of compound **15**. These data showed again that the shape of the 3-substituent is an important factor, which is responsible for the compound’s biological activity. Unfortunately, the steric contribution of the 3-position substituents is difficult to predict due to the absence of data regarding target protein/compound interactions.

**Table 1.**
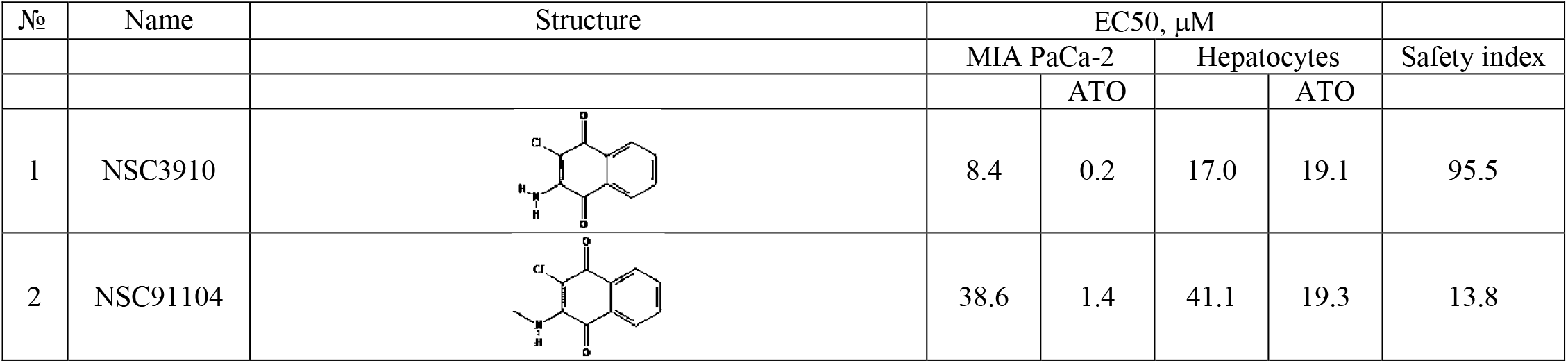
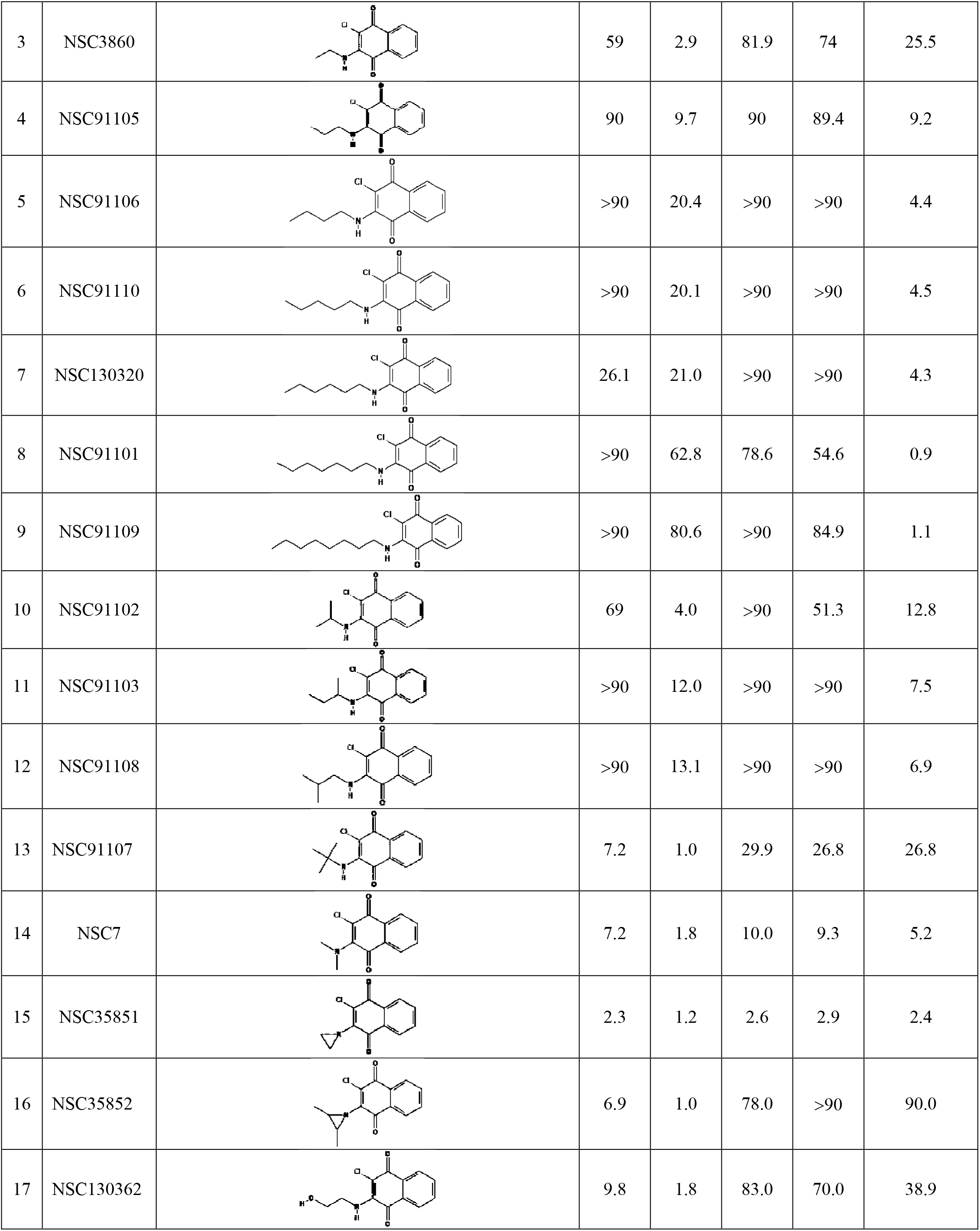
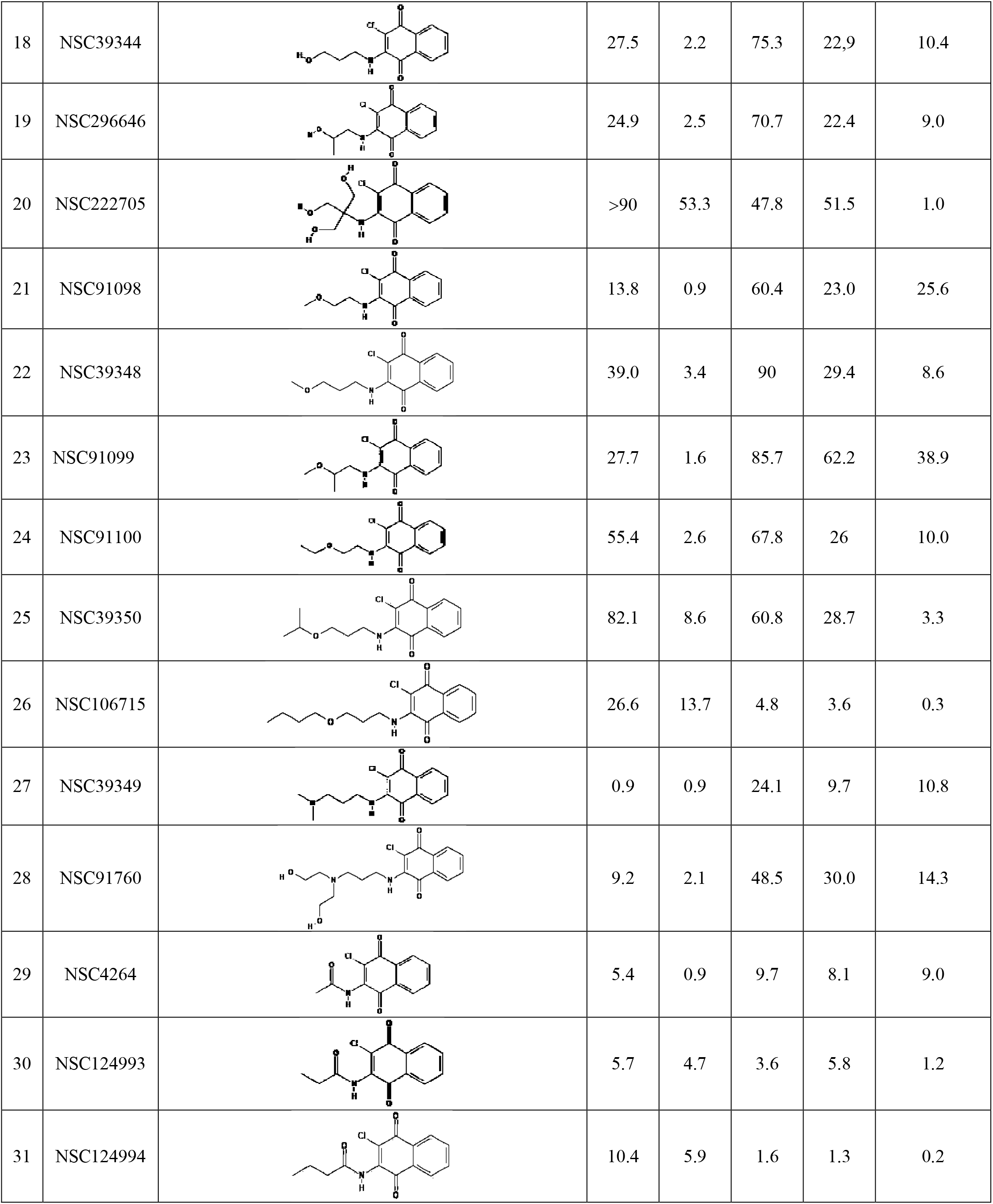
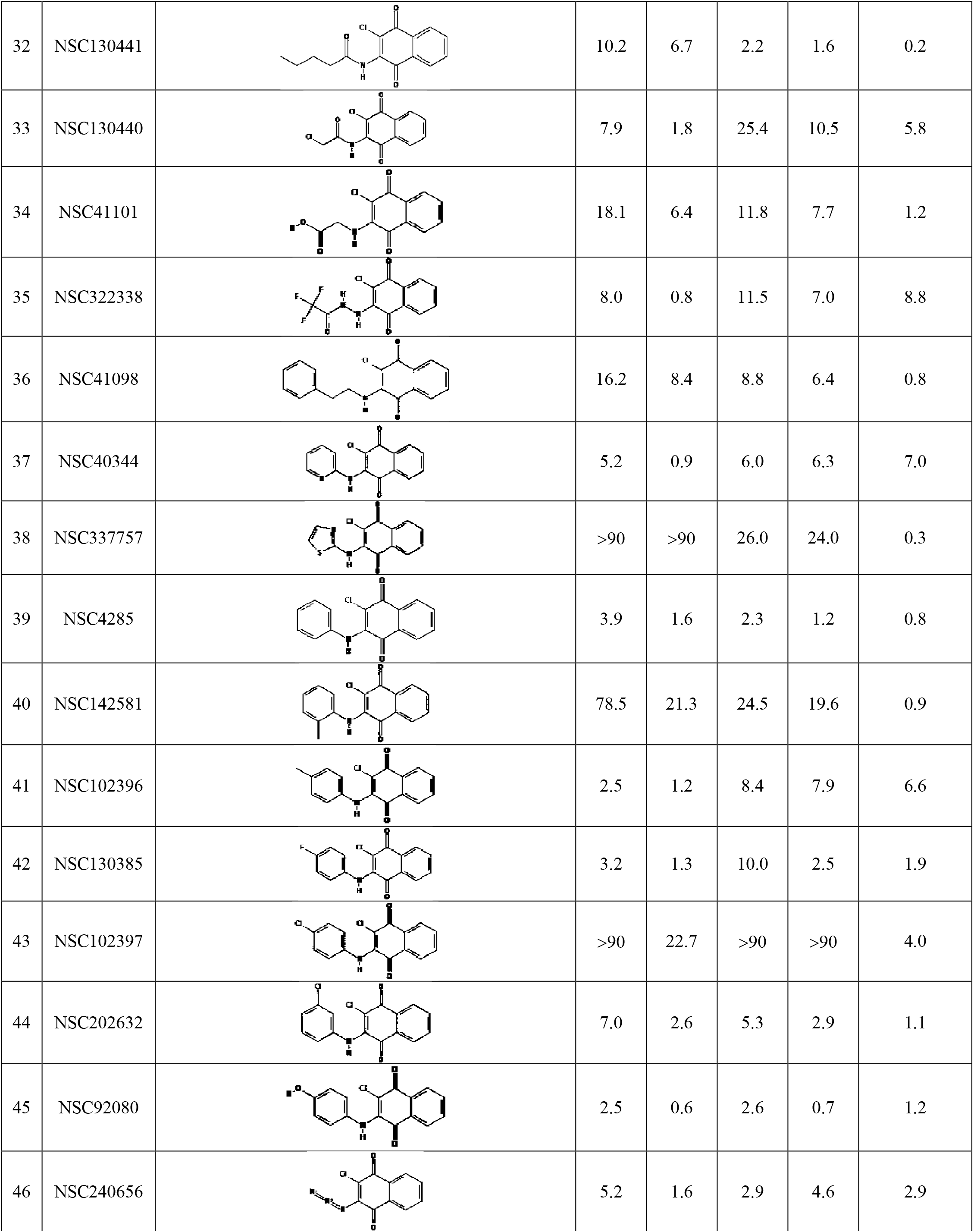
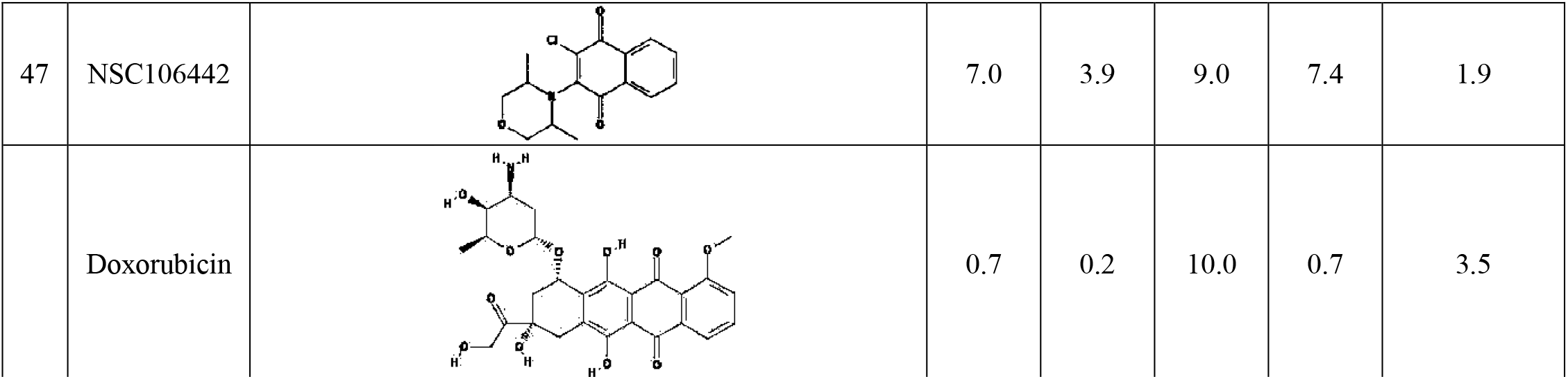
EC50 values and the safety index of the NSC130362 analogs that contain 3-chloro substituent in MIA PaCa-2 cells and hepatocytes. Doxorubicin was included as a reference compound. Standard deviation for all EC50 values does not exceed 10%. To calculate the safety index for compounds that have the EC50 value above 90 μM, EC50 values were equated to 90 μM. *P*<0.05.

| № | Name | Structure | EC50, μM |  |  |  | Safety index |
| --- | --- | --- | --- | --- | --- | --- | --- |
|  |  |  | MIA PaCa-2 |  | Hepatocytes |  |  |
|  |  |  |  | ATO |  | ATO |  |
| 1 | NSC3910 |  | 8.4 | 0.2 | 17.0 | 19.1 | 95.5 |
| 2 | NSC91104 |  | 38.6 | 1.4 | 41.1 | 19.3 | 13.8 |
| 3 | NSC3860 |  | 59 | 2.9 | 81.9 | 74 | 25.5 |
| 4 | NSC91105 |  | 90 | 9.7 | 90 | 89.4 | 9.2 |
| 5 | NSC91106 |  | >90 | 20.4 | >90 | >90 | 4.4 |
| 6 | NSC91110 |  | >90 | 20.1 | >90 | >90 | 4.5 |
| 7 | NSC130320 |  | 26.1 | 21.0 | >90 | >90 | 4.3 |
| 8 | NSC91101 |  | >90 | 62.8 | 78.6 | 54.6 | 0.9 |
| 9 | NSC91109 |  | >90 | 80.6 | >90 | 84.9 | 1.1 |
| 10 | NSC91102 |  | 69 | 4.0 | >90 | 51.3 | 12.8 |
| 11 | NSC91103 |  | >90 | 12.0 | >90 | >90 | 7.5 |
| 12 | NSC91108 |  | >90 | 13.1 | >90 | >90 | 6.9 |
| 13 | NSC91107 |  | 7.2 | 1.0 | 29.9 | 26.8 | 26.8 |
| 14 | NSC7 |  | 7.2 | 1.8 | 10.0 | 9.3 | 5.2 |
| 15 | NSC35851 |  | 2.3 | 1.2 | 2.6 | 2.9 | 2.4 |
| 16 | NSC35852 |  | 6.9 | 1.0 | 78.0 | >90 | 90.0 |
| 17 | NSC130362 |  | 9.8 | 1.8 | 83.0 | 70.0 | 38.9 |
| 18 | NSC39344 |  | 27.5 | 2.2 | 75.3 | 22,9 | 10.4 |
| 19 | NSC296646 |  | 24.9 | 2.5 | 70.7 | 22.4 | 9.0 |
| 20 | NSC222705 |  | >90 | 53.3 | 47.8 | 51.5 | 1.0 |
| 21 | NSC91098 |  | 13.8 | 0.9 | 60.4 | 23.0 | 25.6 |
| 22 | NSC39348 |  | 39.0 | 3.4 | 90 | 29.4 | 8.6 |
| 23 | NSC91099 |  | 27.7 | 1.6 | 85.7 | 62.2 | 38.9 |
| 24 | NSC91100 |  | 55.4 | 2.6 | 67.8 | 26 | 10.0 |
| 25 | NSC39350 |  | 82.1 | 8.6 | 60.8 | 28.7 | 3.3 |
| 26 | NSC106715 |  | 26.6 | 13.7 | 4.8 | 3.6 | 0.3 |
| 27 | NSC39349 |  | 0.9 | 0.9 | 24.1 | 9.7 | 10.8 |
| 28 | NSC91760 |  | 9.2 | 2.1 | 48.5 | 30.0 | 14.3 |
| 29 | NSC4264 |  | 5.4 | 0.9 | 9.7 | 8.1 | 9.0 |
| 30 | NSC124993 |  | 5.7 | 4.7 | 3.6 | 5.8 | 1.2 |
| 31 | NSC124994 |  | 10.4 | 5.9 | 1.6 | 1.3 | 0.2 |
| 32 | NSC130441 |  | 10.2 | 6.7 | 2.2 | 1.6 | 0.2 |
| 33 | NSC130440 |  | 7.9 | 1.8 | 25.4 | 10.5 | 5.8 |
| 34 | NSC41101 |  | 18.1 | 6.4 | 11.8 | 7.7 | 1.2 |
| 35 | NSC322338 |  | 8.0 | 0.8 | 11.5 | 7.0 | 8.8 |
| 36 | NSC41098 |  | 16.2 | 8.4 | 8.8 | 6.4 | 0.8 |
| 37 | NSC40344 |  | 5.2 | 0.9 | 6.0 | 6.3 | 7.0 |
| 38 | NSC337757 |  | >90 | >90 | 26.0 | 24.0 | 0.3 |
| 39 | NSC4285 |  | 3.9 | 1.6 | 2.3 | 1.2 | 0.8 |
| 40 | NSC142581 |  | 78.5 | 21.3 | 24.5 | 19.6 | 0.9 |
| 41 | NSC102396 |  | 2.5 | 1.2 | 8.4 | 7.9 | 6.6 |
| 42 | NSC130385 |  | 3.2 | 1.3 | 10.0 | 2.5 | 1.9 |
| 43 | NSC102397 |  | >90 | 22.7 | >90 | >90 | 4.0 |
| 44 | NSC202632 |  | 7.0 | 2.6 | 5.3 | 2.9 | 1.1 |
| 45 | NSC92080 |  | 2.5 | 0.6 | 2.6 | 0.7 | 1.2 |
| 46 | NSC240656 |  | 5.2 | 1.6 | 2.9 | 4.6 | 2.9 |
| 47 | NSC106442 |  | 7.0 | 3.9 | 9.0 | 7.4 | 1.9 |
|  | Doxorubicin |  | 0.7 | 0.2 | 10.0 | 0.7 | 3.5 |

Next, we analyzed different 2-chloro-3-(hydroxylalkylamino)-1,4-NQs where the alkyl group contains single hydroxyl groups (compounds **17–19**). In contrast to compounds **3** and **4**, change of the 2-hydroxyethyl group (compound **17**) for either 3-hydroxypropyl (compound **18**) or 2-hydroxypropyl (compound **19**) had little effect on the anti-cancer activity but increased toxicity to hepatocytes approximately 3-fold. In compounds **3** and **4**, changing of the ethyl group for the propyl group had little effect on toxicity to hepatocytes but reduced anticancer activity more than 3-fold. On the other hand, exchange of the ethyl group in compound **3** for the isopropyl group (compound **10)**, had little effect on activity against cancer cells or hepatocytes. We conclude again that shape of the alkyl chain in the 3-position contributes to the biological activities of the compounds. To analyze the effect of hydroxyl substitution in the alkyl group, we compared compounds **3** *vs* **17**, and **4** *vs* **18** *vs* **19**. While, compounds **3** and **17** had similar anti-cancer potency and toxicity towards hepatocytes, addition of a hydroxyl group to the propyl group to generate 3-hydroxypropylamino (compound **18**) and 2-hydroxypropylamino (compound **19**) substituent resulted in approximately 2- and 4-fold increase in anti-cancer activity and toxicity to hepatocytes, respectively. We conclude that the hydroxyl group non-specifically increases cytotoxic activity of compounds **18** and **19**. In contrast, addition of 3 hydroxyl groups to the tert-butylamino substituent of compound **13** to generate compound **20**, resulted in approximately 50- and 2-fold loss in anti-cancer activity and toxicity to hepatocytes, respectively. These data suggest that the effect of hydroxyl group in the 3-alkylamino substituent is variable and depends on the length and shape of the alkyl chain.

We also analyzed the effect of different alkoxyamino substituents on biological activity of 2-chloro-3- (alkoxyamino)-1,4-NQs (compounds **21–26**). As in the case with compounds **3** and **4**, change of the 2-methoxyethylamino substituent (compound **21**) for 3-methoxypropylamino group (compound **22**) resulted in a 4-fold decrease in anti-cancer activity with little effect on toxicity to hepatocytes. However, the effect of attachment of a methoxy group to the C-2 position of the propylamino substituent (compound **23**) on anti-cancer activity is less pronounced with a decrease in toxicity to hepatocytes of almost 3-fold. These data again point to the significance of steric effects of the 3-position substituents in 2-chloro-3-(alkoxyamino)-1,4-NQs. A similar trend in decreased anti-cancer activity with increasing the length of 3-substituent was seen when the 2-methoxyethylamino substituent in compound **21** was changed for the 2-ethoxyethylamino (compound **24**), the 3-isopropoxypropyl (compound **25**) or the 3-butoxypropylamino group (compound **26**). However, while compounds **21**, **24**, and **25** have similar toxicity to hepatocytes (EC50 = 23–29), the presence of 3-butoxypropylamino substituent in compound **26** drastically increased toxicity to hepatocytes (EC50 = 3.6). The positive affect of a methoxy group on the anti-cancer activity was seen by comparing compounds **3** *vs* **21**, and **4** *vs* **22** but the presence of a methoxy group also increased toxicity to hepatocytes approximately 3-fold. In contrast, attachment of a methoxy group at the C-2 position of 3-propylamino substituent (compound **23**) resulted in a 6-fold improved anti-cancer activity without significant effect on toxicity to hepatocytes as compared to compound **4**. By comparing compounds **21** *vs* **24**, and **22** *vs* **25** *vs* **26,** it was evident that increasing the length of alkoxy attachment to the 3-alkylamino substituent decreased anti-cancer activity and can, as in the case of compound **26**, increase toxicity to hepatocytes. We conclude that because the alkoxy group can have different effects on anticancer activity and toxicity to hepatocytes (compare compounds **4** *vs* **22** and **4** *vs* **23**), it is possible, *via* careful selection of the C-3 position substituent, to improve anti-cancer potency while maintaining low toxicity to hepatocytes. A 10.8-fold improvement in the anti-cancer activity of compound **4** was seen when the 3-propylamino substituent was extended by the dimethylamino group (compound **27**), however, it also increased toxicity to hepatocytes 9.2 times. When the 3-propylamino substituent of compound **4** was extended by the bis(2-hydroxyethyl)amino moiety (compound **28**), anti-cancer activity and toxicity to hepatocytes were increased 4.6- and 3.0-fold, respectively.

Both, the presence of carbonyl group and increasing the length of the C-3 position substituent were unfavorable in relation to the development goals as was evidenced by gradual decrease in the anti-cancer activity and an increase in the toxicity to hepatocytes of compounds **29**, **30**, **31**, and **32** as compared to compound **1**. Substitution of hydrogen with a chloro group in the 3-acethylamnio substituent of compound **29** to generate compound **33**, caused 2-fold decrease in the anti-cancer activity. Extension of the C-3 position amino substituent in compound **1** by either the 2-oxo-2-hydroxyethyl (compound **34**) or the 2,2,2-trifluoroacetamide group (compound **35**) also decreased the anti-cancer activity and increased toxicity to hepatocytes. We conclude that in all cases the presence of a carbonyl group in the C-3 position substituent made the compound less potent in relation to cancer cells and more toxic in relation to normal cells indicating a loss of selectivity toward cancer cells. Similarly, either a decrease in anti-cancer activity and/or loss of tumor selectivity was observed when the extension of the 3-amino substituent in compound **1** was a phenethyl (compound **36**),a pyridine (compound **37**), a 1,3-thiazole (compound **38**) or a phenyl group with or without fluorine, chlorine, hydroxyl, and methyl groups at different positions (compounds **39–45**). We conclude that various extensions of the 3-amino group of 2-chloro-3-amino-1,4-NQ is detrimental to anti-cancer activity and safety index. A lack of significant cancer cell specificity and reduced anticancer activity were also observed regardless of whether triazadien or 3,5-dimethylmorpholin was at the C-3 position (compounds **46** and **47**). The highest safety index (>90) and low EC50 (0.2 and 1.0 μM) in cancer cells were for compounds that have a 3-amino (compound **1**) and a 3-(2,3-dimethylaziridin-1-yl) (compound **16**) substituent, respectively. For comparison, a potent anti-cancer drug doxorubicin showed similar EC50 in MIA PaCa-2 cells (0.2 μM), but its safety index was approximately 25 times less (Table 1).

According to our MS/MS analysis, NSC130362 efficiently reacts with GSH and most likely with any accessible sulfhydryl group present in a cysteine residue of proteins [8]. We believe that this reactivity is responsible for the short half-life of NSC130362 in the bloodstream. In the previous studies, we also showed that this reaction does not affect the ability of NSC130362 to inhibit GSR and cause oxidative stress. Based on this information, we selected the next set of 1,4-NQs that do not contain 2-chloro substituent (Table 2).

**Table 2.**
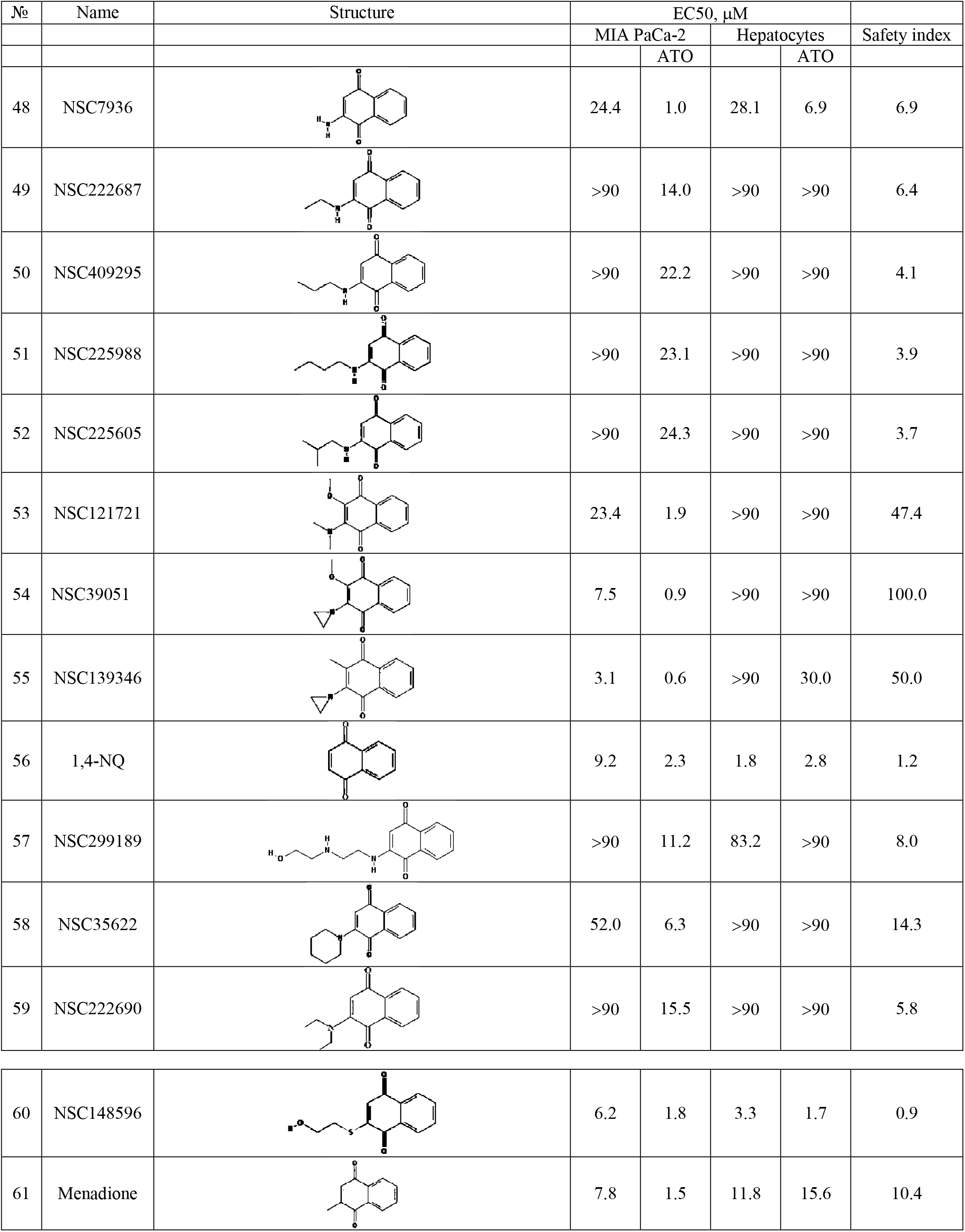
EC50 values and the safety index of the NSC130362 analogs that do not contain 2-chloro substituent in MIA PaCa-2 cells and hepatocytes. Standard deviation for all EC50 values does not exceed 10%. To calculate the safety index for compounds that have the LC50 value above 90 μΜ, EC50 values were equated to 90 μΜ. *P*<0.05.

| № | Name | Structure | EC50, μM |  |  |  | Safety index |
| --- | --- | --- | --- | --- | --- | --- | --- |
|  |  |  | MIA PaCa-2 |  | Hepatocytes |  |  |
|  |  |  |  | ATO |  | ATO |  |
| 48 | NSC7936 |  | 24.4 | 1.0 | 28.1 | 6.9 | 6.9 |
| 49 | NSC222687 |  | >90 | 14.0 | >90 | >90 | 6.4 |
| 50 | NSC409295 |  | >90 | 22.2 | >90 | >90 | 4.1 |
| 51 | NSC225988 |  | >90 | 23.1 | >90 | >90 | 3.9 |
| 52 | NSC225605 |  | >90 | 24.3 | >90 | >90 | 3.7 |
| 53 | NSC121721 |  | 23.4 | 1.9 | >90 | >90 | 47.4 |
| 54 | NSC39051 |  | 7.5 | 0.9 | >90 | >90 | 100.0 |
| 55 | NSC139346 |  | 3.1 | 0.6 | >90 | 30.0 | 50.0 |
| 56 | 1,4-NQ |  | 9.2 | 2.3 | 1.8 | 2.8 | 1.2 |
| 57 | NSC299189 |  | >90 | 11.2 | 83.2 | >90 | 8.0 |
| 58 | NSC35622 |  | 52.0 | 6.3 | >90 | >90 | 14.3 |
| 59 | NSC222690 |  | >90 | 15.5 | >90 | >90 | 5.8 |
| 60 | NSC148596 |  | 6.2 | 1.8 | 3.3 | 1.7 | 0.9 |
| 61 | Menadione |  | 7.8 | 1.5 | 11.8 | 15.6 | 10.4 |

We compared 1,4-NQs with 2-chloro-1,4-NQs that have identical C-3 position substituent to determine the role of the 2-chloro group in compound activity. Removal of the C-2 position chloro group in compounds **3**, **4**, **5**, and **12** to generate compounds **48**, **49**, **50**, **51**, and **52** reduced anti-cancer activity. When the C-3 position substituent was a primary amino group (compounds **1** and **48**), this chlorine removal increased toxicity to hepatocytes approximately 3-fold. In all other pairs, removal of the C-2 chloro group did not significantly affect toxicity to hepatocytes. We also noticed that with increasing the length of 3-alkylamino substituent, the effect of 2-chloro group on the anti-cancer potency was decreased. For example, when compounds have a 3-ethylamino substituent (compounds **3** and **49**), a 2-chlorine insertion was associated with a marked increase in cytotoxicity in MIA PaCa-2 cells, while the presence of a 3-propylamino (compounds **4** and **50**), 3-isobutylamino (compounds **12** and **52**), and 3-butylamino (compounds **5** and **51**) substituent gradually decreased the effect of the 2-position chloro group on anti-cancer activity.

On the other hand, exchange of the C-2 position chlorine in 2-chloro-3-(dimethylamino)-1,4-NQ (compound **14)** and 2-chloro-3-(aziridin-1-yl)-1,4-NQ (compound **15)** for a methoxy group to generate compounds **53** and **54**, respectively did not significantly affect the anti-cancer activity but greatly reduced toxicity to hepatocytes. Moreover, exchange of the C-2 position chlorine in compound **15** for a methyl group (compound **55**) increased the anti-cancer activity 2-fold and reduced toxicity to hepatocytes more than 10-fold. Again, we see the opposite effect of the substitution on the activity toward cancer and normal cells. Based on these data, we conclude that the C-2 position chlorine substituent (such as in NSC130362) was not necessary for the anti-cancer activity and when a compound contains either the C-3 position dimethylamino (compound **53**) or aziridin-1-yl (compound **54** and **55**) substituent, it resembles NSC130362 in terms of anti-cancer activity and low level of toxicity to hepatocytes.

We also analyzed 1,4-NQs with or without C-3 position amino substituents. In comparing 1,4-NQ (compound **56**) with 3-amino-1,4-NQ (compound **48**) and 3-alkylamino-1,4-NQs (compounds **49–52**), it was found that, in contrast to compounds **48–52**, 1,4-NQ does not exhibit any tumor selectivity. Among compounds **48–52** and **56**, 3-amino-1,4-NQ (compound **48**) was the most active against cancer cells and its tumor selectivity was reflected by the safety index equal to 6.9. In contrast to 1,4-NQ and 3-amino-1,4-NQ (compounds **56** and **48**), all tested 3-alkylamino-1,4-NQs (compounds **49–52**) were completely non-toxic to human hepatocytes (EC50 > 90 μΜ). We conclude that the presence of 3-alkylamino substituent confers tumor selectivity. Similarly, insertion of a 2-(2-hydroxyethylamino)ethylamino, a 1-piperidinyl or a 2-diethylamino group at 3-position (compounds **57**–**59**) increased EC50 in cancer cells compared to that of 1,4-NQ (compound **56**), and eliminated any detectable toxicity to hepatocytes (EC50 > 90 μΜ). In contrast, insertion of a C-3 position 2-hydroxyethylsulfanyl group (compound **60**) resulted in slightly decreased EC50 in both cancer (EC50 = 1.8 μΜ) and normal (EC50 = 1.7 μΜ) cells. Another reference compound, menadione [30] (compound **61**), which is 3-methyl-1,4-NQ, showed 1.5-fold improved anti-cancer activity as compared to 1,4-NQ (compound **56**) and it was approximately 5 times less toxic to hepatocytes (EC50 = 15.6 μΜ).

Finally, we analyzed quinoline-5,6-diones (Table 3). When the naphthalene unit of the 1,4-NQ_backbone of compound **1** (2-chloro-3-amino-1,4-NQ) was replaced with a quinoline, the resulting compound **62** almost completely lost tumor selectivity without losing anti-cancer activity. On the other hand, the same exchange of the 1,4-backbone in compound **14** and **48** to generate compound **63** and **64**, respectively did not significantly affect either the anti-cancer activity or toxicity to hepatocytes, while the conversion of compound **49** into compound **66** improved the anti-cancer activity more than 30 times without affecting toxicity to hepatocytes. These data suggest that either the type of the 3-position substituent or the presence of the 2-chloro group in a 1,4-NQ derivative influences the effect of the naphthalene/quinoline exchange on either the anti-cancer activity or toxicity to hepatocytes. Furthermore, either complete removal of the C-2 position substituent or exchange of the 2-chloro group with bromine in compound **61** to generate compounds **64** or **65**, reduced both the anti-cancer activity and toxicity to hepatocytes. We conclude that the 7-position substituent in quinoline-5,6-diones, which corresponds to the C-2 position substituent in 1,4-NQs, also contributes to the level of the anti-cancer activity and toxicity to hepatocytes. Interestingly, when the 6-position primary amino group of compound **64** was replaced by an anilino group, creating compound **67**, a well-known GSR inhibitor LY83583, anti-cancer activity increased approximately 10-fold, while toxicity to hepatocytes was decreased 3-fold. It is noteworthy that the effect of the amine/aniline exchange in compound **64** to generate compound **66** is opposite to that of the amine/aniline exchange in compound **1**, where the resulting compound **39** had 8-fold reduced anti-cancer activity and completely lost tumor selectivity. These data again suggest that careful selection of substituents in the quinone unit of either 1,4-NQs or quinoline-5,6-diones can result in improved anti-cancer activity and without negative effect on toxicity to hepatocytes.

**Table 3.**
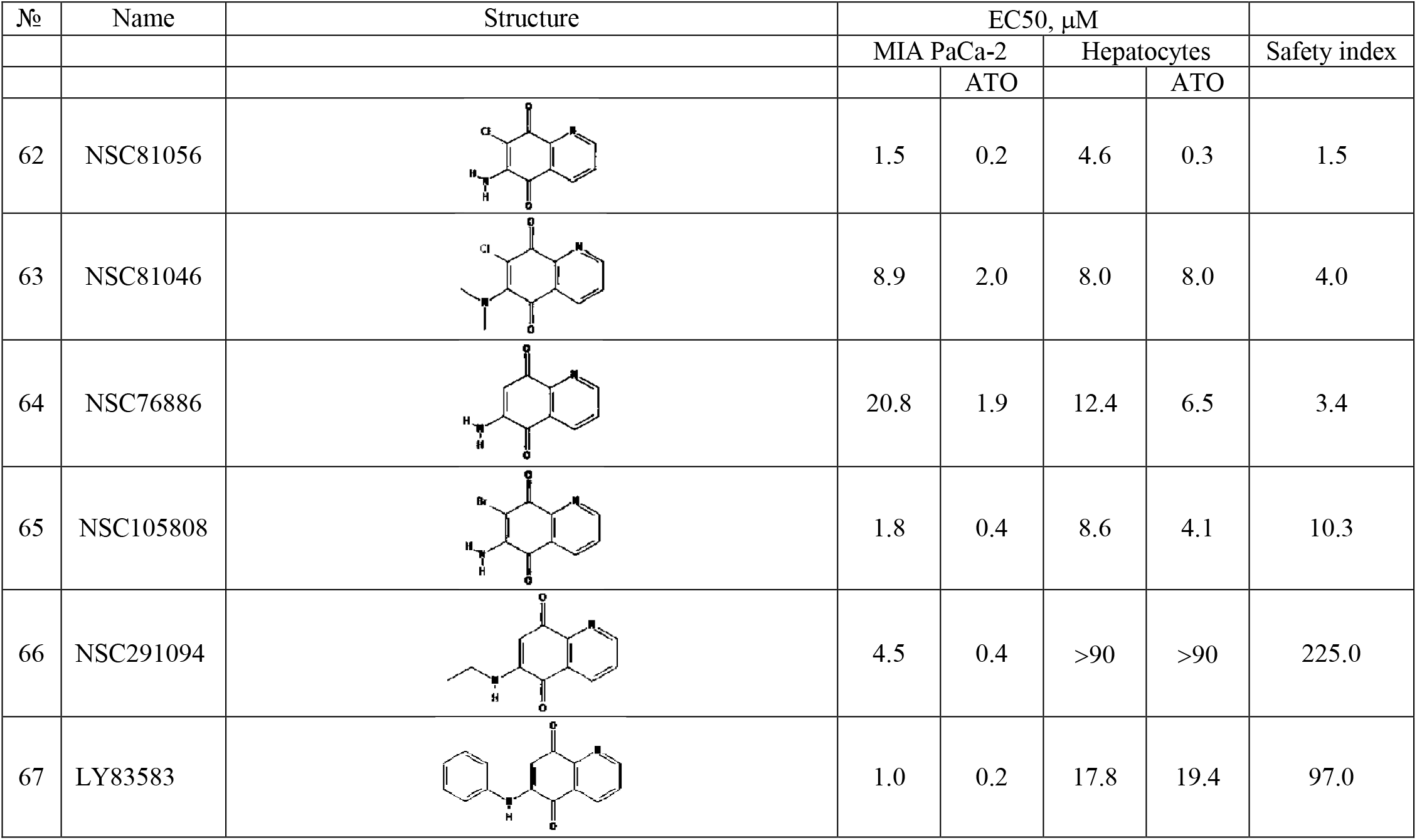
EC50 values and the safety index of NSC130362 analogs in MIA PaCa-2 cells and hepatocytes. Standard deviation for all EC50 values does not exceed 10%. *P*<0.05.

| № | Name | Structure | EC50, μM |  |  |  | Safety index |
| --- | --- | --- | --- | --- | --- | --- | --- |
|  |  |  | MIA PaCa-2 |  | Hepatocytes |  |  |
|  |  |  |  | ATO |  | ATO |  |
| 62 | NSC81056 |  | 1.5 | 0.2 | 4.6 | 0.3 | 1.5 |
| 63 | NSC81046 |  | 8.9 | 2.0 | 8.0 | 8.0 | 4.0 |
| 64 | NSC76886 |  | 20.8 | 1.9 | 12.4 | 6.5 | 3.4 |
| 65 | NSC105808 |  | 1.8 | 0.4 | 8.6 | 4.1 | 10.3 |
| 66 | NSC291094 |  | 4.5 | 0.4 | >90 | >90 | 225.0 |
| 67 | LY83583 |  | 1.0 | 0.2 | 17.8 | 19.4 | 97.0 |

### Activities of the selected NSC130362 analogs

For further analyses, we used compounds that have the safety index either above 20 (non-toxic compounds) or below 1 (toxic compounds) with the aim to identify targets that are responsible for the selective activity of the identified compounds toward cancer cells. To confirm that the effects of the selected compounds are not only specific to MIA PaCa-2 cells, we evaluated their cytotoxic activities on breast carcinoma HCC1187 cells (Table 4). HCC1187 cells were chosen because these cells are highly responsive to ROS [8]. For easy comparison, the data with MIA PaCa-2 cells were also included in Table 4. With only a few exceptions, all non-toxic compounds had a safety index above 10 in HCC1187 cells. We also noticed that, again with a few exceptions, HCC1187 cells are more resistant to the non-toxic compounds in the presence of ATO as compared to MIA PaCa-2 cells. In contrast, HCC1187 cells are more sensitive to all toxic compounds in the presence of ATO as compared to MIA PaCa-2 cells. The high sensitivity of HCC1187 cells to toxic compounds contributed to the increased safety index. In fact, all selected compounds had a safety index above 1 for HCC1187 cells. Based on these results we conclude that despite the difference in sensitivity to some compounds, targeting both HCC1187 and MIA Paca-2 cells by the non-toxic compounds most likely occurs *via* shared mechanisms.

**Table 4.** Activities of the selected NSC130362 analogs. ^a^ Enzyme residual activity after treatment as compared to DMSO-treated sample. ^b^ Concentration of H_2_O_2_ (μM) generated in NADH/NADPH-dehydrogenase-mediated reaction. *P*<0.05.

| № | Name | LC50 |  | LC50 |  | LC50 |  | Safety index |  | ROS increase, % |  | Mito<br>Check<br>I<sup>a</sup> | GSR<sup>a</sup> | NADH/<br>NADPH<br>dehydro<br>genase<sup>b</sup> | GAP<br>DH<sup>a</sup> | LogD,<br>pH =<br>7.4 |
| --- | --- | --- | --- | --- | --- | --- | --- | --- | --- | --- | --- | --- | --- | --- | --- | --- |
|  |  | HCC1187 |  | MIA PaCa-2 |  | Hepatocytes |  | HCC<br>1187 | MIA<br>PaCa-2 | MIA<br>PaCa-2 | Hepato<br>-cytes |  |  |  |  |  |
|  |  |  | ATO |  | ATO |  | ATO | ATO | ATO |  |  |  |  |  |  |  |
| 66 | NSC291094 | 13.2 | 4.4 | 4.5 | 0.4 | >90 | >90 | 20.5 | 225.0 | 626 | 173 | 158 | 59 | 2.87 | 93 | 2.87 |
| 54 | NSC39051 | 18.2 | 2.2 | 7.5 | 0.9 | >90 | >90 | 40.9 | 100.0 | 675 | 124 | 84 | 54 | 0.31 | 73 | 0.31 |
| 67 | LY83583 | 3.0 | 0.2 | 1.0 | 0.2 | 17.8 | 19.4 | 97.0 | 97.0 | 783 | 179 | 116 | 53 | 3.45 | 76 | 3.45 |
| 1 | NSC3910 | 10.8 | 2.2 | 8.4 | 0.2 | 17.0 | 19.1 | 8.7 | 95.5 | 331 | 156 | 145 | 33 | 2.01 | 100 | 2.01 |
| 16 | NSC35852 | 3.6 | 0.7 | 6.9 | 1.0 | 78.0 | >90 | 128.6 | 90.0 | 637 | 180 | 58 | 75 | 0.70 | 100 | 0.70 |
| 55 | NSC139346 | 51.8 | 1.9 | 3.1 | 0.6 | >90 | 30.0 | 15.8 | 50.0 | 416 | 125 | 72 | 83 | 0.12 | 100 | 0.12 |
| 53 | NSC121721 | >90 | 9.5 | 23.4 | 1.9 | >90 | >90 | 9.5 | 47.4 | 446 | 117 | 51 | 61 | 0.00 | 95 | 0.00 |
| 17 | NSC130362 | 19.2 | 1.5 | 9.8 | 1.8 | 83.0 | 70.0 | 46.7 | 38.9 | 295 | 132 | 38 | 65 | 1.12 | 89 | 1.12 |
| 23 | NSC91099 | 19.2 | 5.0 | 27.7 | 1.6 | 85.7 | 62.2 | 12.4 | 38.9 | 284 | 133 | 93 | 66 | 0.86 | 100 | 0.86 |
| 13 | NSC91107 | 28.7 | 1.5 | 7.2 | 1.0 | 29.9 | 26.8 | 17.9 | 26.8 | 248 | 123 | 84 | 45 | 0.52 | 100 | 0.52 |
| 12 | NSC91098 | 5.2 | 4.7 | 13.8 | 0.9 | 60.4 | 23.0 | 4.9 | 26.8 | 254 | 154 | 104 | 65 | 0.00 | 100 | 0.00 |
| 3 | NSC3860 | 23.6 | 4.4 | 59 | 2.9 | 81.9 | 74 | 16.8 | 25.5 | 182 | 115 | 93 | 59 | 1.01 | 100 | 1.01 |
| 60 | NSC148596 | 0.6 | 0.2 | 6.2 | 1.8 | 3.3 | 1.7 | 8.5 | 0.9 | 2188 | 386 | 212 | 38 | 2.99 | 88 | 2.99 |
| 40 | NSC142581 | 9.0 | 5.1 | 78.5 | 21.3 | 24.5 | 19.6 | 3.8 | 0.9 | 111 | 122 | 103 | 72 | 0.00 | 88 | 0.00 |
| 8 | NSC91101 | 56.1 | 22.3 | >90 | 62.8 | 78.6 | 54.6 | 2.4 | 0.9 | 110 | 115 | 116 | 93 | 0.17 | 100 | 0.17 |
| 36 | NSC41098 | 2.6 | 2.6 | 16.2 | 8.4 | 8.8 | 6.4 | 2.5 | 0.8 | 721 | 178 | 135 | 42 | 0.00 | 97 | 0.00 |
| 39 | NSC4285 | 1.7 | 0.4 | 3.9 | 1.6 | 2.3 | 1.2 | 3.0 | 0.8 | 1247 | 174 | 91 | 92 | 0.13 | 93 | 0.13 |
| 38 | NSC337757 | 20.3 | 8.1 | >90 | >90 | 26.0 | 24.0 | 3.0 | 0.3 | 100 | 100 | 105 | 90 | 0.10 | 100 | 0.10 |
| 26 | NSC106715 | 1.7 | 0.7 | 26.6 | 13.7 | 4.8 | 3.6 | 5.1 | 0.3 | 889 | 191 | 112 | 66 | 0.00 | 94 | 0.00 |
| 32 | NSC130441 | 0.3 | 0.7 | 10.2 | 6.7 | 2.2 | 1.6 | 2.3 | 0.2 | 506 | 249 | 135 | 1 | 2.79 | 3 | 2.79 |
| 31 | NSC124994 | 1.0 | 0.5 | 10.4 | 5.9 | 1.6 | 1.3 | 2.6 | 0.2 | 251 | 228 | 159 | 28 | 2.03 | 5 | 2.03 |

The marked difference in the anti-tumor activity and safety index between the non-toxic and toxic compounds (Table 4) indicates that these two groups of compounds should differ in either physicochemical properties or biological activities. Analysis of lipophilicity indicates that both the non-toxic and toxic compounds have similar distribution coefficient LogD that ranges from 0 to 3.45. We conclude that the analysis of the octanol-water partitioning is not sufficient to establish a link between degree of lipophilicity and biological activities of the selected compounds.

The cytotoxic effects of quinones have been attributed to several mechanisms: (i) DNA damage mediated *via* inhibition of DNA topoisomerase-II [31]; (ii) production of ROS *via* NADH/NADPH dehydrogenase-mediated formation of semiquinone radical that can reduce oxygen to produce super oxide [16]; (iii) inhibition of glyceraldehyde 3-phosphate dehydrogenase (GAPDH) [17], (iv) inhibition of GSR [8, 17, 32], (v) alkylation of cellular nucleophiles such as GSH, DNA, RNA and proteins, and (vi) inhibition of mitochondrial function [17, 21]. To elucidate the type of compound activity which is responsible for its cytotoxic effects in cancer cells and which is tolerated in normal cells, we performed various *in vitro* and cell-based activity assays (Table 4).

#### ROS induction in MIA Paca-2 cells and hepatocytes

As the first step, we evaluated the ability of the selected compounds to induce ROS in cancer and normal cells. Cells were incubated with the compounds for 4 h and the level of ROS induction was measured by a hydrogen peroxide cell-based assay kit. Compounds were assayed at a fixed concentration of 30 μM and the measured level of hydrogen peroxide was expressed as percentage in relation to DMSO-treated cells (100%). As shown in Table 4, all non-toxic compounds induced more ROS in cancer cells than in hepatocytes. This distinction was more evident for compounds with the highest safety index. Approximately half of the toxic compounds also induced more ROS in cancer cells than in normal cells. We conclude that the selective induction of ROS in cancer cells is required but not sufficient factor for the selective anti-tumor activity.

#### NADH/NADPH dehydrogenase-mediated redox cycling

As the next step, we examined the capability of the selected compounds to generate ROS in NADH/NADPH dehydrogenase-mediated reaction *in vitro* (Table 4). ROS can be generated by NADH/NADPH-dehydrogenase *via* the one electron reduction of a quinone ring of 1,4-NQs [16, 33]. Unstable half-reduced quinones (semiquinones) transfer electrons to molecular oxygen, thus generating superoxide anion radicals (O_2_^•−^), which are converted by SOD to hydrogen peroxide [34]. To determine the ability of compounds to induce ROS, compounds were incubated for 4 h at 37°C with and without NADH/NADPH-dehydrogenase in the presence of NADPH as an electron donor and SOD to convert generated superoxide radicals into hydrogen peroxide. At the end of incubation, 10-acetyl-3,7-dihydroxyphenoxazine (ADHP) and horseradish peroxidase (HRP) were added to the reaction and the level of HRP-mediated formation of the fluorogenic product, resorufin, was measured on a plate reader. Except for a few compounds, ROS production was detected in reactions with either the toxic or non-toxic compounds. Again, both the non-toxic and toxic compounds could either generate H_2_O_2_ by NADH/NADPH-dehydrogenase-mediated reaction or be inert in this activity. Based on these results, we conclude that the ability to generate ROS *via* NADH/NADPH-dehydrogenase mediated redox cycling cannot be the sole factor contributing to the selective anti-cancer activity of the tested compounds.

#### Effect on mitochondrial function

To further investigate the mechanism of anti-cancer activity, we evaluated the ability of the selected compounds to target mitochondria because inhibition of mitochondrial function by 1,4-NQs can be lethal to a cell [17]. Complex I consisting of NADH dehydrogenase constitutes one of the major sites of electron entry into the mitochondrial electron transport chain (ETC). Complex I catalyzes the transfer of two electrons from NADH to ubiquinone to form ubiquinol and finally to the terminal electron acceptor, molecular oxygen. During this process, four protons are moved from mitochondria to the transmembrane space forming a pH gradient and an electrical potential across the inner mitochondrial membrane, which are required for oxidative phosphorylation. To test possible effects of compounds on ETC, we analyzed the Complex I-mediated NADH oxidation using a MitoCheck Complex I Activity Assay Kit. The level of NADH oxidation was measured by a decrease in absorbance at 340 nm and expressed as percentage of residual activity in relation to DMSO-treated mitochondria (Table 4). Addition of either non-toxic or toxic compounds to bovine heart mitochondria at a concentration of 30 μM caused both inhibition and stimulation of NADH oxidation. We noticed that the set of the non-toxic compounds mostly consists of inhibitors of Complex I activity, while the set of the toxic compounds does not have any Complex I inhibitory activity, except compound **39** that inhibited mitochondrial function by 9%. It is noteworthy that because both Complex I and NADH/NADPH-dehydrogenase use NADH as a source of electrons, observed activity of Complex I does not correspond to the actual Complex I activity. Complex I activity was estimated by the rate of NADH oxidation which was also affected by NADH/NADPH-dehydrogenase-mediated redox cycling. Indeed, we noticed that compounds **1**, **31**, **32**, **60**, and **65** that stimulated Complex I Activity by more than 30% also induced a high level of ROS in the NADH/NADPH dehydrogenase-mediated reaction *in vitro.* Thus, increased activity of Complex I most likely was observed due to the increased NADH oxidation by NADH/NADPH-dehydrogenase. On the other hand, compounds that both inhibited Complex I and stimulated ROS production actually were more potent inhibitors than can be estimated by the level of Complex I inhibition, because they also promoted NADH oxidation by NADH/NADPH-dehydrogenase. Altogether, these data suggest that there was a bias of the non-toxic compounds to inhibitors of Complex I activity.

#### GSR Inhibition activity

Next, we evaluated the inhibitory activity of the selected compounds against human GSR enzyme using recombinant GSR and Abcam GSH/GSSG ratio detection reagent (Table 4). Briefly, GSR activity was measured by following oxidized glutathione (GSSG) reduction in the presence of NADPH and expressed as percentage of residual activity with respect to DMSO-treated enzyme. The residual activity of GSR for both the non-toxic and toxic compounds was quite heterogeneous, ranging from 100% (compounds **38**) to 3% (compound **32**). Remarkably, two compounds **31** and **32** with the lowest safety index were the most potent inhibitors of GSR. We thus conclude that GSR inhibitory activity alone cannot be used to predict the safety index.

#### GAPDH Inhibition Activity

As a final step, we evaluated the inhibitory activity of the selected compounds against human GAPDH enzyme in a cell-based assay using MIA PaCa-2 cells (Table 4). Cells were pre-incubated with compounds for 2 h at 37°C followed by cell lysis using sonication and GAPDH activity assay. The ability of compounds to inhibit GAPDH was expressed as percentage of GAPDH residual activity with respect to DMSO-treated cells. As shown in Table 4, most compounds inhibited GAPDH by 0 to 27%. Similar to the GSR activity assay, only two of the most toxic compounds **31** and **32**, were able to inhibit GAPDH by 95 and 97%, respectively. Based on these data, we conclude that inhibition of GAPDH is not the activity responsible for the selective cytotoxicity of the non-toxic compounds.

### Quantitative SAR (QSAR) model

The mechanism of anti-tumor action of the non-toxic compounds should differ from that of the toxic compounds. Based on the obtained results we determined that except targeting mitochondrial ETC, the non-toxic compounds do not exhibit biological activities that are clearly distinct from those of toxic compounds. Since targeting mitochondria could involve different proteins and enzymes and chemical reactions of quinones are complex, we assumed that molecular modeling, which takes into account all available physicochemical properties of analyzed molecules, is an appropriate tool to identify key structural features that are associated with the observed selective cytotoxic activity. To accomplish this task, we employed the Q-MOL molecular modeling package (www.q-mol.com) [8, 35, 36] that uses a novel proprietary approach to take into account molecular flexibility, a common roadblock for many other molecular modeling tools. Taking into account the structure of the non-toxic and toxic compounds, Q-MOL developed a mathematical model or fitness function to distinguish accurately compounds with desired properties (anti-tumor activity and safety toward normal cells) (Fig 6). This fitness function was based on atom field potentials (AFPs) which, in turn, were derived from molecular mechanics force field. As shown in Fig 6, the resulting mathematical model shows that the non-toxic compounds have higher (less negative) AFPs than the toxic compounds.

**Fig 6.**
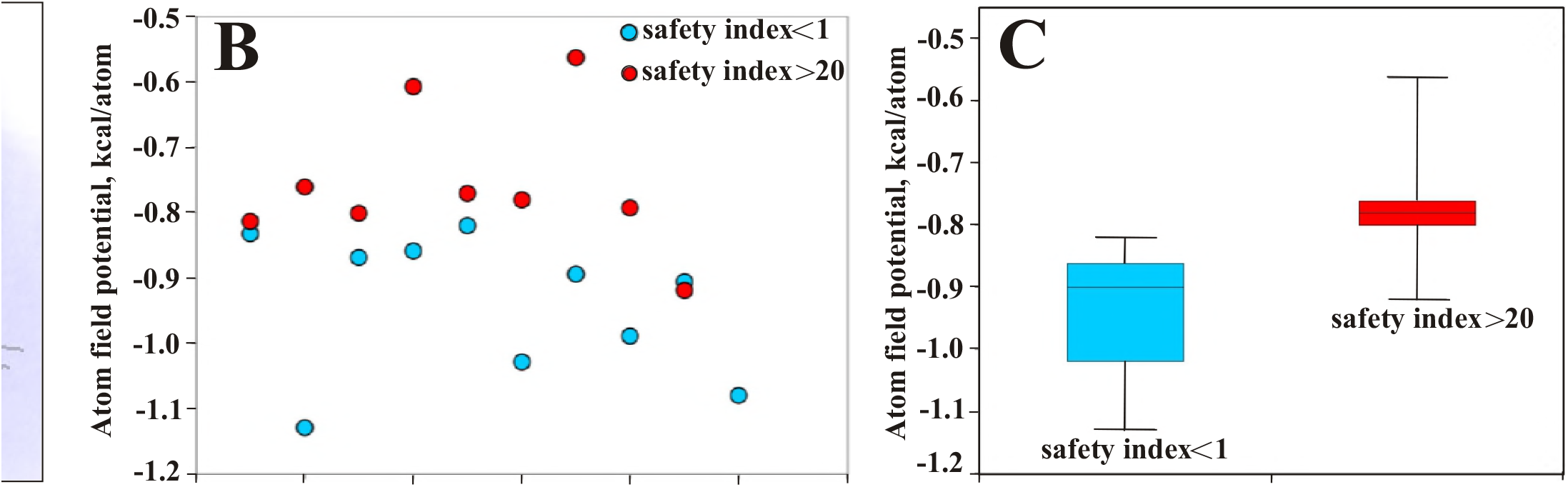
QSAR modeling of NSC130362 analogs. *A.* 3-D alignment of NSC130362 analogs. Red, blue and green color denote oxygen, nitrogen, and chlorine atoms, respectively; *B,* the distribution of the non-toxic (safety index above 20) and toxic (safety index below 1) analogs in relation to AFP; *C*, an average of AFPs for the nontoxic and toxic compounds. *P*<0.05.

## Discussion

Recently, we published that NSC130362 inhibits GSR and, as a consequence, induces ROS and subsequent apoptosis in cancer cells but not in normal human hepatocytes as a model for hepatotoxicity [8]. The goal of this study was two-fold: [1] to identify specific biological activities or structural features, which are responsible for the observed selective anti-cancer activity and [2] to determine whether a QSAR model could be designed for the development of potent anti-cancer compounds that are non-toxic toward normal cells. The NIH has defined several necessary conditions that should be met for validation of a compound to be target for optimization studies. Some of these conditions such as reproducible response in different assay types and adequate anti-tumor activity were addressed in our previous work [8]. To further validate NSC130362 as a candidate molecule for further characterization and/or development, we performed additional cell-based assays. ROS inducers and GSR inhibitors are also considered as drug sensitizers [22–24, 37]. Indeed, we showed that NSC130362 potentiated anti-tumor potency of all tested pancreatic and prostate cancer drugs and decreased EC50 (cytotoxicity) of several kinase inhibitors by up to 3-logs (1000 times) in leukemia cells directly *ex vivo* from patients. The ability to potentiate the cytotoxic activity of various anti-cancer agents is a valuable property of NSC130362. This conclusion is based on the fact that under treatment, cancer cells often adapt and develop resistance to anti-cancer drugs. To maximally exploit chemotherapy as a therapeutic strategy, it would be meaningful to combine anticancer agents with compounds that target these adaptive mechanisms. One of the mechanisms of adaptation that can confer chemoresistance is upregulation of antioxidant pathways. In agreement, there is accumulating evidence showing improved treatment outcome when anti-cancer drugs are combined with ROS-inducing agents [22–24]. In line with this evidence, our data also suggest that anti-cancer activity of the NSC130362/ATO combination is affected by the reducing environment inside cancer cells. It may be the case that targeting GSR activity, which controls the reducing environment in cancer cells, makes them susceptible to ATO and thus inducing apoptosis by altering redox-sensitive proteins and enzymes [27]. The reason the effect of hypoxic conditions on the activity of the NSC130362/ATO combination is not pronounced is likely because ETC can function efficiently at oxygen levels as low as 0.5% [38].

In addition, NSC130362 belongs to the class of 1,4-NQs that consists of important and widely distributed compounds with a variety of pharmacological properties. 1,4-NQs contain two ketone groups which have the ability to accept electrons and generate ROS *via* NADH/NADPH dehydrogenase-mediated redox cycling mechanism [16, 33]. 1,4-NQs can also react with different biological targets. The NCI have identified the quinone moiety as an important pharmacophore element for anti-cancer drugs [39] because it possesses several tumor-selective activities such as anti-proliferative and anti-angiogenesis activity [40, 41], ability to induce oxidative stress, apoptosis and necrosis [42], ability to inhibit DNA replication [43], ability to suppress glycolysis and mitochondrial function [16], and ability to inhibit NF-κB and STAT3 signaling [37, 44]. As a consequence, quinone moieties are present in many cancer drugs such as anthracyclines, daunorubicin, doxorubicin, and mitomycin [16].

To determine structural features of NSC130362, which are responsible for the selective anti-cancer activity, we performed SAR analysis of NSC130362 analogs. In the course of SAR studies, we focused on changes to the quinone moiety of NSC130362 because 1,4-NQ derivatives with various substitutions in C-2 and C-3 positions have been shown to exhibit potent anti-cancer properties [20, 45]. Because we were interested in identifying mechanisms underlying the selective anti-cancer cytotoxicity, the non-toxic compounds (safety index above 20] and toxic compounds (safety index below 1] were, therefore, selected for further experiments. Our studies support the following model: (1) With a few exceptions, non-toxic compounds exhibited lower lipophilicity as compared to toxic compounds. These results are in agreement with recent studies showing that the most potent anti-tumor compound had the highest hydrophilicity (lowest lipophilicity) among a aminonaphthoquinone series [46]. (2) One of the most important findings is that the C-2 chloro group of NSC130362 is dispensable for its activity and it can be removed to potentially improve pharmacokinetics (PK) and safety of the compound. The highly reactive C-2 chloro group may contribute to the short half-life of NSC130362 in the mouse bloodstream [8]. In addition, in agreement with our data, chlorine insertion at C-3 contributes to redox potential and reduced the safety index, which was determined by comparing the IC50 measured in cancer cells versus that obtained in normal fibroblasts [46]. These results are consistent with the chloro group at the C-2 position contributing to selectivity toward cancer cells. (3) The presence of the carbonyl group in the C-3 substitution group in all cases reduced the safety index, indicating a loss of selectivity toward cancer cells. (4) Two most toxic analogs (MIA PaCa-2-related safety index = 0.2) almost completely inhibited GAPDH. Previous papers described the involvement of GAPDH in growth and programmed cell death in hepatocytes suggesting a link between GAPDH and toxicity to hepatocytes [47, 48]. However, further analysis is needed to determine whether the GAPDH inhibitory activity is associated with toxicity in hepatocytes. The findings indicate a strong SAR among different 1,4-NQs and quinoline-5,8- diones and highlight the significance of functional groups as substituents in these compounds affecting their anticancer activity and safety index.

1,4-NQ has limited solubility in aqueous solutions. Converting the naphthalene backbone of 1,4-NQ into a quinoline by introduction of a nitrogen atom at C-5 or at C-8 can increase aqueous solubility. However, introduction of a nitrogen atom at the C-5 or at C-8 of 1,4-NQ derivative menadione can increase the oxidant character of the molecule [32, 49]. To explore this modification, we compared compounds **1** *vs* **62**, **14** *vs* **63**, **48** *vs* **64**, and **49** *vs* **66**. While in the case of compounds **1** *vs* **62**, the introduction of quinoline resulted in 64-fold increase in toxicity to hepatocytes without any noticeable effect on anti-cancer activity, the same modification in either compound **14** or **48** did not significantly affect either anti-cancer activity or toxicity to hepatocytes. In contrast, conversion of 1,4-NQ backbone in compound **49** to generate compound **66** resulted in more than 30fold improved anti-cancer activity without any visible effect on toxicity to hepatocytes. We conclude that the effect of naphthalene/quionoline exchange depends on the presence of the 2-chloro substituent and functional groups at the C-3 and, most likely, at other positions. More detailed analysis of different substitutions in the described or other positions of either naphthalene or quinolone backbones would require additional compounds be synthesized to make conclusive observations which is beyond the scope of this study.

With the aim of identifying the biological activity of the tested compounds that is responsible for the selective cytotoxicity in cancer cells, we tested the non-toxic (safety index is above 20) and toxic (safety index is below 1) compounds in a variety of cell-based and *in vitro* assays. In earlier studies it has been shown that the cytotoxic activity of 1,4-NQ is directly dependent on the redox potential, which accounts for the semiquinone production followed by autoxidation of semiquinone and generation of ROS [16, 20]. To analyze the ability to generate ROS, we tested the selected compounds in cell-based assays as well as in direct *in vitro* ROS assays using recombinant NADH/NADPH dehydrogenase. While all non-toxic compounds preferentially induce ROS in cancer cells, five out of nine toxic compounds also induced more ROS in cancer cells than in normal cells. In addition, the results of NADH/NADPH dehydrogenase-based *in vitro* assay confirmed that ROS-inducing capability cannot be the sole factor contributing to the value of the safety index. Moreover, we noticed that three toxic compounds did not generate any ROS in NADH/NADPH dehydrogenase-mediated reaction. We conclude that at least some toxic compounds induce ROS in cancer cells and hepatocytes by a mechanism that does not involve NADH/NADPH dehydrogenase.

To control the level of ROS and maintain redox homeostasis, mammalian cells have developed multiple strategies that inactivate ROS and prevent its accumulation. One such strategy is governed by the GSH/GSR-based electron donor system. Thus, compounds that target GSR can induce oxidative stress and subsequent cell death. To explore this possibility, we tested the selected compounds in *in vitro* GSR activity assays. All non-toxic compounds exhibited a moderate level of GSR-inhibiting activity ranging from 17% (compound **55**) to 67% (compound **1**), while the toxic compounds had much wider range from 7% (compound **8**) to 99% (compound **32**). Most likely, compounds **26**, **36**, and **40**, which did not produce ROS by NADH/NADPH dehydrogenase *in vitro,* induced ROS in cancer and hepatocytes by inhibiting GSR. We also noticed that the level of induced ROS in cancer cells does not necessarily correlate with the ability of compounds to produce ROS *in vitro* and inhibit GSR. This discrepancy can be explained by the different ability of compounds to penetrate cells where the compound’s intracellular concentration could be 100–1000 times higher than its extracellular concentration as has been described for phenylethyl isothiocyanate (PEITC) [50, 51]. In addition, there are additional pathways that regulate ROS production and levels in cells that do not involve GSR. These could be manifest in intact cells.

The effect of 1,4-NQs on mitochondrial function [17] could play a major role in their activity and selectivity. Indeed, we found that most non-toxic compounds inhibited oxygen consumption supported by Complex I in bovine heart mitochondria. It is noteworthy that mitochondrial respiration was stimulated by the Complex I substrate NADH, which is also used as a donor of electrons in NADH/NADPH dehydrogenase-mediated reduction of 1,4-NQs. Mitochondrial function was measured by the rate of NADH oxidation. Thus, actual inhibition of mitochondrial function should be greater than that we measured because of these two simultaneous reactions: Complex I and NADH/NADPH dehydrogenase. Indeed, the non-toxic compounds that showed induction of mitochondrial function (compounds **1**, **66**, and **67**) had the highest ability to generate ROS in NADH/NADPH dehydrogenase-mediated reaction. Only one non-toxic compound (compound **12**) was not able both to generate ROS by NADH/NADPH dehydrogenase and to inhibit mitochondrial function. However, this compound efficiently inhibited GSR. Most likely, the cytotoxic effect of compound **12**, was mediated *via* inhibition of GSR.

In contrast, all toxic compounds with one exception (compound **39** with 9% inhibition) did not inhibit mitochondrial respiration. Again, toxic compounds **31**, **32**, and **60** with the highest ROS-inducing ability showed increased NADH oxidation in the Complex I activity assay. Based on these data, we conclude that effects on mitochondrial respiration could contribute to the selective anti-cancer activity of the non-toxic compounds. On the other hand, the toxic compounds induced cytotoxicity by mechanisms that are distinct from inhibition of mitochondrial respiration.

The mitochondria play a critical role in the production of ATP and the metabolites that are necessary for biosynthesis of macromolecules such as nucleotides, fatty acids, and glutathione [52]. The generation of ATP occurs *via* a set of mitochondrial intermembrane complexes I, III, and IV that make up the ETC. Recent studies provided evidence that the mitochondrial ETC might be a valuable target for cancer therapy [52, 53]. For example, it has been shown that metformin exerts anti-tumor effects partially through its ability to inhibit mitochondrial complex I and exhibits an excellent safety profile [52, 54]. Another inhibitor of complex I activity, phenformin, also exerts its anti-tumor effects in orthotopic mouse models [55]. It is noteworthy that, like NSC130362, inhibitors of mitochondrial function, including metformin and phenformin, improve the anti-cancer effects of compounds that target PI3K and B-RAF signaling [56–58]. Mitochondrial complex I inhibitors block tricarboxylic acid (TCA) cycle-mediated ATP production and induce apoptosis by enhancing the generation of ROS. A ROS-based approach to induce selective cytotoxicity in cancer cells has been recently validated by researchers at Kolkata’s Indian Association for the Cultivation of Science (IACS) who have synthesized a novel porphyrin compound with potent anti-cancer activity [59]. By targeting cellular topoisomerase 1 enzyme and producing increased ROS inside cells, this porphyrin compound showed potent cytotoxicity against cervical, ovarian, and colon cancer cells with significantly reduced or no toxicity toward human embryonic kidney cell lines and mouse embryonic fibroblasts.

Although most cancer cells have functional mitochondrial metabolism, some cancer cells have mutations in the ETC and macromolecule biosynthesis proteins of their mitochondria [60, 61]. These cancer cells are dependent on glycolysis to produce energy (Warburg effect). Thus, inhibiting glycolysis is an efficient way to target these ETC-mutated cancer cells. In addition, one of the key hallmarks of cancer is to re-program metabolic capability, which leads to elevated glycolytic metabolism in cancer cell [62]. Thus, it has been hypothesized that inhibition of both mitochondrial metabolism and glycolysis could be an effective anti-cancer therapy [63]. GAPDH catalyzes the conversion of glyceraldehyde 3-phosphate to D-glycerate 1,3-bisphosphate in the process of glycolysis [64]. Because GAPDH is one of the possible targets of 1,4-NQ, we evaluated the selected compounds in cell-based GAPDH inhibition assays. None of the non-toxic compounds inhibited GAPDH by more than 27%. In contrast, two the most toxic compounds (compounds 3**1** and **32**) almost completely inhibited GAPDH. Because the same compounds were the most efficient inhibitors of GSR, we speculated that this dual effect can be responsible for the high level of toxicity to hepatocytes and, as a consequence, the lowest safety index of compounds **31** and **32**. Other toxic compounds inhibited GAPDH with narrow range of 0–12%. We conclude that targeting glycolysis is not the factor that is responsible of the selective anti-cancer activity of the non-toxic compounds.

The results described above allow us to predict that targeting mitochondrial ETC could be the key biological activity that can be used to guide non-toxic compound optimization. We can also speculate that distinct effects of the non-toxic and toxic compounds on mitochondrial function are governed by multiple structural factors that include but are not limited to electronic, hydrophobic, and steric characteristics of substituents and their position in the quinone ring [65, 66]. To account for available physico-chemical factors that most likely affect compound activities, we performed molecular modeling to develop a QSAR model. Molecular modeling employed computational-based descriptors to correlate biological activity in cellular systems. Based on the structure of the non-toxic and toxic compounds, Q-MOL developed a fitness function (AFPs) to describe accurately the relationship between the molecular structure of a chemical and its anti-tumor potency and safety toward normal cells. Encouragingly, there was a clear difference in the AFPs between the non-toxic and toxic compounds, which confirmed that the AFP-based QSAR model could be trained by docking ligands into AFP distributions and then used as an energy-based classifier to differentiate between non-toxic and toxic ligands. Therefore, we conclude that it is feasible, under appropriate settings, to develop non-toxic compounds with potent anti-tumor activity.

## References

1. Nordhoff A, Bücheler US, Werner D, Schirmer RH. Folding of the four domains and dimerization are impaired by the Gly446-->Glu exchange in human glutathione reductase. Implications for the design of antiparasitic drugs. Biochemistry. 1993; 32: 4060–4066.

2. Sosa V, Moliné T, Somoza R, Paciucci R, Kondoh H, LLeonart ME. Oxidative stress and cancer: an overview. Ageing Res Rev. 2013; 12: 376–390.

3. Reuter S, Gupta SC, Chaturvedi MM, Aggarwal BB. Oxidative stress, inflammation, and cancer: how are they linked? Free Radic Biol Med. 2010; 49: 1603–1616.

4. Pervaiz S, Clement MV. Tumor intracellular redox status and drug resistance--serendipity or a causal relationship? Curr Pharm Des. 2004; 10: 1969–7197.

5. Trachootham D, Zhou Y, Zhang H, Demizu Y, Chen Z, Pelicano H, et al. Selective killing of oncogenically transformed cells through a ROS-mediated mechanism by beta-phenylethyl isothiocyanate. Cancer Cell. 2006; 10: 241–252.

6. Zhang R, Humphreys I, Sahu RP, Shi Y, Srivastava SK. In vitro and in vivo induction of apoptosis by capsaicin in pancreatic cancer cells is mediated through ROS generation and mitochondrial death pathway. Apoptosis. 2008; 13: 1465–1478.

7. Pelicano H, Carney D, Huang P. ROS stress in cancer cells and therapeutic implications. Drug Resist Updat. 2004; 7: 97–110.

8. Rozanov D, Cheltsov A, Sergienko E, Vasile S, Golubkov V, Aleshin AE, et al. TRAIL-based high throughput screening reveals a link between TRAIL-mediated apoptosis and glutathione reductase, a key component of oxidative stress response. PLoS One. 2015; 10: e0129566.

9. Couzin J. Cancer drugs. Smart weapons prove tough to design. Science 2002; 298: 522–525.

10. Frantz S. Drug discovery: playing dirty. Nature. 2005; 437: 942–943.

11. Lu J, Holmgren A. Thioredoxin system in cell death progression. Antioxid Redox Signal. 2012; 17: 1738–1747.

12. Couto N, Wood J, Barber J. The role of glutathione reductase and related enzymes on cellular redox homoeostasis network. Free Radic Biol Med. 2016; 95: 27–42.

13. Kamada K, Goto S, Okunaga T, Ihara Y, Tsuji K, Kawai Y, et al. Nuclear glutathione S-transferase pi prevents apoptosis by reducing the oxidative stress-induced formation of exocyclic DNA products. Free Radic Biol Med. 2004; 37: 1875–1884.

14. Fernando MR, Lechner JM, Löfgren S, Gladyshev VN, Lou MF. Mitochondrial thioltransferase (glutaredoxin 2) has GSH-dependent and thioredoxin reductase-dependent peroxidase activities in vitro and in lens epithelial cells. FASEB J. 2006; 20: 2645–2647.

15. Geisbrecht BV, Gould SJ. The human PICD gene encodes a cytoplasmic and peroxisomal NADP(+)-dependent isocitrate dehydrogenase. J Biol Chem. 1999; 274: 30527–33053.

16. O’Brien PJ. Molecular mechanisms of quinone cytotoxicity. Chem Biol Interact. 1991; 80: 1–41.

17. Prati F, Bergamini C, Molina MT, Falchi F, Cavalli A, Kaiser M, et al. 2-Phenoxy-1,4-naphthoquinones: From a Multitarget Antitrypanosomal to a Potential Antitumor Profile. J Med Chem. 2015; 58: 6422–6434.

18. Yan C, Siegel D, Newsome J, Chilloux A, Moody CJ, Ross D. Antitumor indolequinones induced apoptosis in human pancreatic cancer cells via inhibition of thioredoxin reductase and activation of redox signaling. Mol Pharmacol. 2012; 81: 401–410.

19. Hughes JP, Rees S, Kalindjian SB, Philpott KL. Principles of early drug discovery. Br J Pharmacol. 2011; 162: 1239–1249.

20. Prachayasittikul V, Pingaew R, Worachartcheewan A, Nantasenamat C, Prachayasittikul S, Ruchirawat S, et al. Synthesis, anticancer activity and QSAR study of 1,4-naphthoquinone derivatives. Eur Med Chem. 2014; 84: 247–263.

21. Nicotera P, Hinds TR, Nelson SD, Vincenzi EF. Differential effects of arylating and oxidizing analogs of N-acetyl-p-benzoquinoneimine on red blood cell membrane proteins, Arch. Biochem. Biophys. 1990; 283: 200–205.

22. Maiti AK. Genetic determinants of oxidative stress-mediated sensitization of drug-resistant cancer cells. International Journal of Cancer. 2012; 130: 1–9.

23. Donadelli M, Costanzo C, Beghelli S, Scupoli MT, Dandrea M, Bonora A, et al. Synergistic inhibition of pancreatic adenocarcinoma cell growth by trichostatin A and gemcitabine. Biochimica et Biophysica Acta. 2007; 1773: 1095–1106.

24. Maeda H, Hori S, Ohizumi H, Segawa T, Kakehi Y, Ogawa O, et al. Effective treatment of advanced solid tumors by the combination of arsenic trioxide and L-buthionine-sulfoximine. Cell Death and Differentiation. 2004; 11: 737–746.

25. Hidalgo M. Pancreatic Cancer. N Engl J Med. 2010; 362: 1605–1617.

26. Tyner JW, Yang WF, Bankhead A 3rd, Fan G, Fletcher LB, Bryant J, et al. Kinase pathway dependence in primary human leukemias determined by rapid inhibitor screening. Cancer Res. 2013; 73: 285–296.

27. Miller WH Jr, Schipper HM, Lee JS, Singer J, Waxman S. Mechanisms of action of arsenic trioxide. Cancer Res. 2002; 62: 3893–3903.

28. Ramadori G, Cameron S. Effects of systemic chemotherapy on the liver. Ann Hepatol. 2010; 9: 133–143.

29. Zhou S, Palmeira CM, Wallace KB. Doxorubicin-induced persistent oxidative stress to cardiac myocytes. Toxicol Lett. 2001; 121: 151–157.

30. Criddle DN, Gillies S, Baumgartner-Wilson HK, Jaffar M, Chinje EC, Passmore S, et al. Menadione-induced reactive oxygen species generation via redox cycling promotes apoptosis of murine pancreatic acinar cells. J Biol Chem. 2006; 281: 40485–40492.

31. Bender RP, Ham AJ, Osheroff N. Quinone-induced enhancement of DNA cleavage by human topoisomerase IIalpha: adduction of cysteine residues 392 and 405. Biochemistry. 2007; 46: 2856–2864.

32. Morin C, Besset T, Moutet JC, Fayolle M, Brückner M, Limosin D, et al. The aza-analogues of 1,4-naphthoquinones are potent substrates and inhibitors of plasmodial thioredoxin and glutathione reductases and of human erythrocyte glutathione reductase. Org Biomol Chem. 2008; 6: 2731–2742.

33. Buffinton GD, Ollinger K, Brunmark A, Cadenas E. DT-diaphorase-catalysed reduction of 1,4-naphthoquinone derivatives and glutathionyl-quinone conjugates. Effect of substituents on autoxidation rates. Biochem J. 1989; 257: 561–571.

34. McCord JM, Fridovich I. Superoxide dismutase: the first twenty years (1968-1988). Free Radic Biol Med. 1988; 5: 363–369.

35. Chen F, Chen J, Lin J, Cheltsov AV, Xu L, Chen Y, et al. NSC-640358 acts as RXRα ligand to promote TNFα-mediated apoptosis of cancer cell. Protein Cell. 2015; 6: 654–666.

36. Shiryaev SA, Cheltsov AV, Gawlik K, Ratnikov BI, Strongin AY. Virtual ligand screening of the National Cancer Institute (NCI) compound library leads to the allosteric inhibitory scaffolds of the West Nile Virus NS3 proteinase. Assay Drug Dev Technol. 2011; 9: 69–78.

37. Sandur SK, Ichikawa H, Sethi G, Ahn KS, Aggarwal BB. Plumbagin (5-hydroxy-2-methyl-1,4-naphthoquinone) suppresses NF-kappaB activation and NF-kappaB-regulated gene products through modulation of p65 and IkappaBalpha kinase activation, leading to potentiation of apoptosis induced by cytokine and chemotherapeutic agents. J Biol Chem. 2006; 281: 17023–17033.

38. Rumsey WL, Schlosser C, Nuutinen EM, Robiolio M, Wilson DF. Cellular energetics and the oxygen dependence of respiration in cardiac myocytes isolated from adult rat. J Biol Chem. 1990; 265: 15392–15402.

39. Liu KC, Li J, Sakya S. Synthetic approaches to the 2003 new drugs. Mini-Rev Med Chem. 2004; 4: 1105–1125.

40. Hafeez BB, Zhong W, Fischer JW, Mustafa A, Shi X, Meske L, et al. Plumbagin, a medicinal plant (Plumbago zeylanica)-derived 1,4-naphthoquinone, inhibits growth and metastasis of human prostate cancer PC-3M-luciferase cells in an orthotopic xenograft mouse model. Mol Oncol. 2013; 7: 428–439.

41. Kayashima T, Mori M, Yoshida H, Mizushina Y, Matsubara K. 1,4-Naphthoquinone is a potent inhibitor of human cancer cell growth and angiogenesis. Cancer Lett. 2009; 278: 34–40.

42. Han W, Li L, Qiu S, Lu Q, Pan Q, Gu Y, et al. Shikonin circumvents cancer drug resistance by induction of a necroptotic death. Mol Cancer Ther. 2007; 6: 1641–1649.

43. Shahabuddin MS, Gopal M. Genotoxicity of DNA Intercalating Anticancer Drugs: Pyrimido{4(I),5(I):4,5} thieno(2,3-b)quinolines on Somatic and Germinal Cells. Toxicol Mech Methods. 2007; 17: 135–145.

44. Bhasin D, Etter JP, Chettiar SN, Mok M, Li PK. Antiproliferative activities and SAR studies of substituted anthraquinones and 1,4-naphthoquinones. Bioorg Med Chem Lett. 2013; 23: 6864–2867.

45. Sunassee SN, Veale CG, Shunmoogam-Gounden N, Osoniyi O, Hendricks DT, Caira MR, et al. Cytotoxicity of lapachol, β-lapachone and related synthetic 1,4-naphthoquinones against oesophageal cancer cells. Eur J Med Chem. 2013; 62: 98–110.

46. Benites J, Valderrama JA, Bettega K, Pedrosa RC, Calderon PB, Verrax J. Biological evaluation of donor-acceptor aminonaphthoquinones as antitumor agents. Eur J Med Chem. 45 2010; 45: 6052–6057.

47. Barbini L, Rodríguez J, Dominguez F, Vega F. Glyceraldehyde-3-phosphate dehydrogenase exerts different biologic activities in apoptotic and proliferating hepatocytes according to its subcellular localization. Mol Cell Biochem. 2007 300: 19–28.

48. Ciriaci N, Vigo B, Rigalli J, Mottino A, Ruiz M, Ghanem C. Acetaminophen (APAP) hepatocyte toxicity is associated with increased translocation of glyceraldehyde-3-phosphate dehydrogenase (GAPDH) to the nucleus. FASEB J. 2015; 29: Suppl 937.8.

49. Shaikh IA, Johnson F, Grollman AP. Streptonigrin. 1. Structure-activity relationships among simple bicyclic analogues. Rate dependence of DNA degradation on quinone reduction potential. J Med Chem. 1986; 29: 1329–40.

50. Xu K, Thornalley PJ. Involvement of glutathione metabolism in the cytotoxicity of the phenethyl isothiocyanate and its cysteine conjugate to human leukaemia cells in vitro. Biochem Pharmacol. 2001; 61: 165–177.

51. Trachootham D, Zhou Y, Zhang H, Demizu Y, Chen Z, Pelicano H, et al. Selective killing of oncogenically transformed cells through a ROS-mediated mechanism by beta-phenylethyl isothiocyanate. Cancer Cell. 2006; 10: 241–252.

52. Weinberg SE, Chandel NS. Targeting mitochondria metabolism for cancer therapy. Nat Chem Biol. 2015; 11: 9–15.

53. Kalyanaraman B, Cheng G, Hardy M, Ouari O, Lopez M, Joseph J, et al. A review of the basics of mitochondrial bioenergetics, metabolism, and related signaling pathways in cancer cells: Therapeutic targeting of tumor mitochondria with lipophilic cationic compounds. Redox Biol. 2018; 14: 316–327.

54. Janzer A, German NJ, Gonzalez-Herrera KN, Asara JM, Haigis MC, Struhl K. Metformin and phenformin deplete tricarboxylic acid cycle and glycolytic intermediates during cell transformation and NTPs in cancer stem cells. Proc Natl Acad Sci U S A. 2014; 111: 10574–10579.

55. Jackson AL, Sun W, Kilgore J, Guo H, Fang Z, Yin Y, et al. Phenformin has anti-tumorigenic effects in human ovarian cancer cells and in an orthotopic mouse model of serous ovarian cancer. Oncotarget. 2017; 8: 100113–100127.

56. Engelman JA, Chen L, Tan X, Crosby K, Guimaraes AR, Upadhyay R, et al. Effective use of PI3K and MEK inhibitors to treat mutant Kras G12D and PIK3CA H1047R murine lung cancers. Nat Med. 2008; 14: 1351–1356.

57. Yuan P, Ito K, Perez-Lorenzo R, Del Guzzo C, Lee JH, Shen CH, et al. Phenformin enhances the therapeutic benefit of BRAF(V600E) inhibition in melanoma. Proc Natl Acad Sci U S A. 2013; 110: 18226–18231.

58. Zheng Z, Zhu W, Yang B, Chai R, Liu T, Li F, et al. The co-treatment of metformin with flavone synergistically induces apoptosis through inhibition of PI3K/AKT pathway in breast cancer cells. Oncol Lett. 2018; 15: 5952–5958.

59. Das SK, Ghosh A, Paul Chowdhuri S, Halder N, Rehman I, Sengupta S, et al. Neutral Porphyrin Derivative Exerts Anticancer Activity by Targeting Cellular Topoisomerase I (Top1) and Promotes Apoptotic Cell Death without Stabilizing Top1-DNA Cleavage Complexes. J Med Chem. 2018; 61: 804–817.

60. Yang M, Soga T, Pollard PJ. Oncometabolites: linking altered metabolism with cancer. J Clin Invest. 2013; 123: 3652–3658.

61. Wallace DC. A mitochondrial paradigm of metabolic and degenerative diseases, aging, and cancer: a dawn for evolutionary medicine. Annu Rev Genet. 2005; 39: 359–407.

62. Ward PS, Thompson CB. Metabolic reprogramming: a cancer hallmark even warburg did not anticipate. Cancer Cell. 2012; 21: 297–308.

63. Cheng G, Zielonka J, Dranka BP, McAllister D, Mackinnon AC Jr, Joseph J, et al. Mitochondria-targeted drugs synergize with 2-deoxyglucose to trigger breast cancer cell death. Cancer Res. 2012; 72: 2634–2644.

64. Ramzan R, Weber P, Linne U, Vogt S. GAPDH: the missing link between glycolysis and mitochondrial oxidative phosphorylation? Biochem Soc Trans. 2013; 41: 1294–1297.

65. Buffinton GD, Ollinger K, Brunmark A, Cadenas E. DT-diaphorase-catalysed reduction of 1,4-naphthoquinone derivatives and glutathionyl-quinone conjugates. Effect of substituents on autoxidation rates. Biochem J. 1989; 257: 561–571.

66. Song Y, Buettner GR. Thermodynamic and kinetic considerations for the reaction of semiquinone radicals to form superoxide and hydrogen peroxide. Free Radic Biol Med. 2010; 49: 919–962.

